# Identification and Characterization of the Lipoprotein *N*-acyltransferase in *Bacteroides*

**DOI:** 10.1101/2024.05.31.596883

**Authors:** Krista M. Armbruster, Jiawen Jiang, Mariana G. Sartorio, Nichollas E. Scott, Jenna M. Peterson, Jonathan Z. Sexton, Mario F. Feldman, Nicole M. Koropatkin

## Abstract

Members of the Bacteroidota compose a large portion of the human gut microbiota, contributing to overall gut health via the degradation of various polysaccharides. This process is facilitated by lipoproteins, globular proteins anchored to the cell surface by a lipidated N-terminal cysteine. Despite their importance, lipoprotein synthesis by these bacteria is understudied. In *E. coli*, the α-amino linked lipid of lipoproteins is added by the lipoprotein *<u>N</u>*-acyltransferase Lnt. Herein, we have identified a protein distinct from Lnt responsible for the same process in *Bacteroides*, named lipoprotein *<u>N</u>*-acyltransferase in *<u>B</u>acteroides* (Lnb). Deletion of Lnb yields cells that synthesize diacylated lipoproteins, with impacts on cell viability and morphology, growth on polysaccharides, and protein composition of membranes and outer membrane vesicles (OMVs). Our results not only challenge the accepted paradigms of lipoprotein biosynthesis in Gram-negative bacteria, but also support the establishment of a new family of lipoprotein *N*-acyltransferases.

**Significance:** Bacteroidota are key members of the human gut microbiota that influence human health by degrading polysaccharides. This degradation is achieved by a suite of lipoproteins, a class of membrane protein characterized by lipidation. Lipoprotein synthesis in Bacteroidota is understudied. Here, we used a genetic screen to identify gene(s) responsible for *N*-acylation, the last step in lipoprotein biosynthesis. Our screen identified the lipoprotein *<u>N</u>*-acyltransferase in *<u>B</u>acteroides* (Lnb) that performs this step. We show that deletion of Lnb negatively affects cellular growth and ability to degrade polysaccharides, deepening our understanding of Bacteroidota and lipoproteins.

## Introduction

Bacteroidota account for nearly 50% of the human gut microbiota and dramatically influence host health and disease (1). These bacteria degrade a wide variety of structurally diverse glycans provided via diet or derived from the host (e.g., intestinal mucin), a metabolic attribute that is key to their survival and persistence in the mammalian gut (2). Breakdown of glycans provides nutrients for fellow gut inhabitants, in turn releasing short-chain fatty acids (SCFAs) that are beneficial to the host (3).

*Bacteroides* spp. devote ∼20% of their genomes to encoding the genes necessary for glycan breakdown in distinct polysaccharide utilization loci (PULs) (4). A typical PUL consists of an outer membrane ß-barrel and one or more associated cell-surface lipoproteins that facilitate capture and breakdown of polysaccharides (5). The most well-studied PUL is the <u>s</u>tarch <u>u</u>ptake <u>s</u>ystem (Sus), comprised of the ß-barrel SusC and four lipoproteins SusDEFG (5). The ability of *Bacteroides* to degrade polysaccharides is intricately linked with lipoprotein biosynthesis and subsequent localization to the cell surface. Many of these lipoproteins are also preferentially packaged onto outer membrane vesicles (OMVs) that have far-reaching impacts on cell and host physiology (6–12). Alongside polysaccharide utilization, lipoproteins are involved in numerous other essential processes, including nutrient uptake, immunomodulation, cell envelope integrity, pilus production, and more (13–16).

Lipoproteins are a class of membrane protein characterized by the post-translational attachment of lipids to a conserved N-terminal cysteine residue. Lipoprotein precursors are initially translated with an N-terminal signal peptide that directs their insertion into the cytoplasmic membrane by the Secretory (Sec) or twin-arginine translocation (Tat) pathways. This signal peptide also encodes a conserved sequence motif, called a lipobox, immediately before the invariant cysteine that marks the precursor for post-translational modification (17). First, a diacylglycerol moiety sourced from a membrane phospholipid is attached to the cysteine thiol by lipoprotein diacyl<u>g</u>lycerol transferase (Lgt) (18). Then, lipoprotein <u>s</u>ignal <u>p</u>eptidase (Lsp) cleaves the signal peptide at the α-amino group of the now-lipidated cysteine (19). In model Gram-negative bacteria such as *Escherichia coli*, lipoprotein *<u>N</u>*-acyltransferase (Lnt) transfers an additional lipid from a second phospholipid donor to form an amide linkage at the cysteine, yielding the mature triacylated lipoprotein (20). *N*-acylated lipoproteins are preferred substrates for the localization <u>o</u>f lipoprotein (Lol) exporter system, which sorts and localizes lipoproteins to the inner or outer membrane based on the amino acids directly following the lipidated cysteine (21–23). Lnt/lipoprotein *N*-acylation is widely regarded as essential to Gram-negative bacteria as a prerequisite to proper lipoprotein localization (22), though there are some known exceptions to this paradigm (24–26).

While Lgt and Lsp are conserved across all bacteria, Lnt is less so, even amongst bacteria known to make triacylated lipoproteins (27, 28). Two such species, *Staphylococcus aureus* and *S. epidermidis*, instead employ the two-component system LnsAB for lipoprotein *N*-acylation (29). *Enterococcus faecalis* and *Bacillus cereus* synthesize lyso-form lipoproteins, with *N*-acylation catalyzed by the lipoprotein intramolecular transacylase Lit (30, 31). Both LnsAB and Lit represent enzyme families unique in sequence and structure to *E. coli* Lnt (29, 32, 33). In fact, all four of these bacterial species were previously presumed to make diacylated lipoproteins based on the absence of an *lnt* ortholog in their genomes. The recent discoveries of Lit and LnsAB demonstrate that assumption of lipoprotein N-terminal structure cannot be made based on the genomic absence of a known lipoprotein modifying enzyme such as Lnt. Furthermore, they also support the possibility of additional lipoprotein biosynthesis pathways that have yet to be identified. Overall, experimental validation of lipoprotein N-terminal structure in various bacteria as well as their corresponding lipoprotein modifying enzymes is necessary to fully understand lipoproteins.

To our knowledge, only one study has empirically characterized lipoprotein N-terminal structure in *Bacteroides*: Hashimoto et al. found that *Bacteroides fragilis* NCTC 9343 synthesizes triacylated lipoproteins (14). However, further investigation into the *B. fragilis* genome by our group reveals no identifiable orthologs of *lnt*. Thus we hypothesized that *B. fragilis* employs distinct enzyme machinery to Lnt to generate triacylated lipoproteins. To identify functional Lnt orthologs in *B. fragilis*, we used a growth rescue strategy involving the *E. coli* conditional lethal *lnt* strain that previously identified Lit as an alternate *N*-acyltransferase (30). With this approach, a library of *B. fragilis* genomic DNA was screened for the ability to functionally rescue Lnt-depleted *E. coli* cells. Herein, we present the results of our experiment, which revealed a single gene responsible for lipoprotein *N*-acylation that is conserved among *Bacteroides* spp. Deletion of this gene negatively affects cell viability and ability to grow on certain polysaccharides, alters cell morphology, and impacts lipoprotein localization and OMV production. Our findings not only demonstrate the importance of lipoprotein *N*-acylation to *Bacteroides* physiology, but also supports the establishment of a new family of lipoprotein *N*-acyltransferases.

## Results

### Identification of *Bacteroides* gene that complements Lnt-depleted *E. coli*

*Bacteroides fragilis* NCTC 9343 produces triacylated lipoproteins (14) despite lacking an ortholog to *E. coli lnt*. In search of candidate Lnt enzymes from *B. fragilis*, we constructed and transformed a library of *B. fragilis* genomic DNA into the *E. coli* conditional-lethal *lnt* strain KA349. In KA349, *lnt* is under the control of the arabinose-inducible promoter *P_BAD_*, thus it requires arabinose for cell survival (30). Transformants were plated onto solid media containing glucose, which represses *P_BAD_*-driven expression of *lnt*. Colonies that grew indicated phenotypic rescue of the lethal depletion of Lnt. Indeed, plasmids encoding two distinct *B. fragilis* genomic inserts were recovered, referred herein as p1 and p31 (Fig. 1A). Both inserts include the open reading frame *BF9343_0945*, a predicted transmembrane protein containing the <u>d</u>omain of <u>u</u>nknown function (DUF) 4105. Plasmid p31 also encodes downstream gene *BF9343_0944*, annotated as an alkaline phosphatase, and the start of *BF9343_0943*, a component of the Sec translocon. Along with *BF9343_0945* and the start of *BF9343_0944*, p1 encodes the pantothenate kinase *BF9343_0946*, putative outer membrane protein *BF9343_0947*, and the start of tetratricopeptide repeat protein *BF9343_0948*.

**Figure 1.**
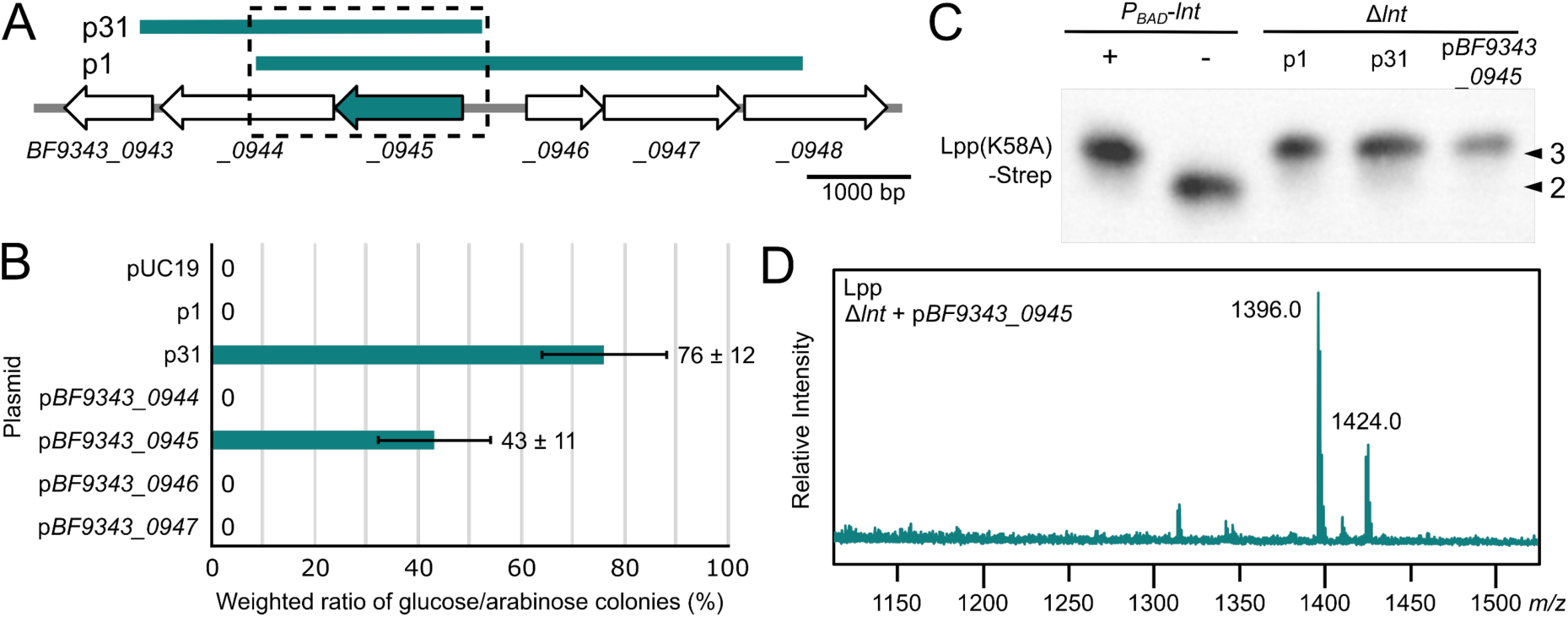
Identification of BF9343_0945 as the lipoprotein triacylating enzyme in Lnt-depleted *E. coli*. (A) DNA fragments recovered from viable Lnt-depleted cells were mapped to the *B. fragilis* genome. The two distinct inserts overlap on the single open reading frame *BF9343_0945* (boxed and shaded). (B) Lnt-depleted KA349 was transformed with the indicated plasmids and spread on three arabinose plates and three glucose plates each. Colonies were enumerated and represented as a weighted ratio of glucose-grown to arabinose-grown colonies. (C) Whole cell lysates of KA349 (*P_BAD_-lnt*) grown under inducing (+) and non-inducing (-) conditions, and Δ*lnt* strains with plasmids p1, p31, and p*BF9343_0945* were separated by SDS-PAGE and immunoblotted against Lpp(K58A)-Strep. Lipoprotein having three acyl chains (arrowhead marked “3”) and two acyl chains (arrowheads marked “2”) are indicated. (D) Trypsinized lipopeptides of Lpp purified from the Δ*lnt* strain expressing BF9343_0945 were eluted from nitrocellulose and analyzed by MALDI-TOF MS. The prominent peak at *m/z* 1396 corresponds to the triacylated N-terminal peptide possessing acyl chains totaling 48:1 (with 48 and 1 referring to the total number of carbons and a double bond, respectively), and *m/z* 1424 is the same peptide with acyl chains totaling 50:1.

To narrow down which gene(s) are responsible for rescue, p1, p31, and plasmids encoding their respective individual genes were tested for the ability to rescue KA349. Lnt-depleted KA349 cells were transformed with each plasmid and spread onto three agar plates containing arabinose and three containing glucose. Regardless of plasmid, all transformations yielded colonies in the presence of arabinose. On glucose, no colonies grew when KA349 was transformed with p*BF9343_0944*, p*BF9343_0946*, and p*BF9343_0947* (Fig. 1B). In contrast, when KA349 was transformed with p31 or p*BF9343_0945*, respectively 76% and 43% of colonies also grew on glucose compared to arabinose.

From these results, we conclude that *BF9343_0945* is sufficient for rescue of KA349, asserting it as our main candidate as the *Bacteroides* lipoprotein *N*-acyltransferase. Despite p1 also encoding *BF9343_0945*, no colonies grew on glucose when KA349 was transformed with p1. This lack of rescue by p1 may be due to the addition of vancomycin to reduce background growth, which was not included in the original rescue experiment.

### BF9343_0945 is sufficient for lipoprotein triacylation in *E. coli*

To demonstrate that the observed rescue cannot be attributed to leaky expression of *P_BAD_-lnt* or residual Lnt enzyme, we sought to determine if *lnt* could be fully deleted in these cells. The marked *lnt::specR* allele from TXM1036, a strain previously used to generate diacylated lipoproteins (30), was packaged into P1*vir* and transduced into Δ*lpp* cells containing p31 and p*BF9343_0945*. We also tested p1 because it rescued KA349 in our initial library screen (Fig. 1A). The *lnt::specR* allele was successfully transduced into cells harboring each of the three plasmids (data not shown), indicating that *lnt* can be deleted when they are present.

We then introduced a plasmid encoding the Strep-tagged lipoprotein Lpp(K58A) into the resulting Δ*lnt* strains and assessed for triacylation by immunoblot. Previously used as a reporter for acylation, the triacylated and diacylated forms of Lpp(K58A)-Strep (a difference of ∼250 Da) can be separated by SDS-PAGE using a 16.5% Tris-tricine gel (30, 34). The triacyl and diacyl controls are derived from KA349 cells grown under Lnt-inducing and Lnt-depleting conditions, respectively. Seen in Fig. 1C, triacylated Lpp was observed in *lnt::specR* cells containing p1, p31, and p*BF9343_0945*. This result not only indicates rescue by lipoprotein triacylation, but also that BF9343_0945 is sufficient for triacylation in *E. coli*.

To further confirm lipoprotein *N*-acylation, we purified Lpp(K58A)-Strep from the Δ*lnt* strain carrying p*BF9343_0945* and analyzed its N-terminal tryptic peptide by matrix-assisted laser desorption ionization-time of flight mass spectrometry (MALDI-TOF MS). As a control, Lpp(K58A)-Strep purified from wildtype (WT) *lnt* cells was also analyzed, whereby a prominent peak appeared at *m/z* 1396 that is consistent with the triacylated-CSSNAK tryptic peptide (Fig. S1A), as previously observed (30). The same peak was detected in the Δ*lnt* strain expressing BF9343_0945 (Fig. 1D). Notably, no peak was observed on either spectrum at *m/z* 1157, the mass corresponding to diacylated-CSSNAK (32). Triacylation was further confirmed in the MS/MS analyses of parent peak *m/z* 1396, revealing the characteristic fragment ions *m/z* 813 and 845 indicative of *N*-acylation (Fig. S1B and S1C)(30).

### Deletion of the candidate Lnt-like protein from *Bacteroides* results in diacylated lipoproteins

We validated our Lnt-like protein in *Bacteroides thetaiotaomicron* (*B. theta*), a close relative of *B. fragilis*. The *B. theta* homolog *BT_4364* is 66% identical (77% similar) to *BF9343_0945* and shares the same genomic neighborhood. Despite the essentiality of Lnt in *E. coli* (22), *BT_4364* was deleted from the *B. theta* genome, albeit with several impacts to cell physiology (described below).

We sought to demonstrate that BT_4364 functions as a lipoprotein *N*-acyltransferase in its native cell background. First, we performed an immunoblot using the same *E. coli* Lpp(K58A)-Strep reporter lipoprotein as in Fig. 1C, recombinantly expressed in *B. theta* from plasmid pWW1376 integrated at an *att* site (35). Lpp(K58A)-Strep from WT *B. theta* cells is consistent in size with the triacylated control from *E. coli*, while Lpp(K58A) from Δ*BT_4364* cells demonstrated a shift in size consistent with the loss of a single acyl chain (Fig. 2A). We then analyzed endogenous *B. theta* lipoprotein BT_3736 (Strep-tagged; chosen for its small size and compatibility with the assay) and observed a similar band shift in Δ*BT_4364* cells (Fig. 2B). In both cases, complementation of the Δ*BT_4364* strain with *BT_4364* under the constitutive promoter *us1311* was sufficient to restore triacylation. Taken together, we conclude that BT_4364 is indeed the lipoprotein *N*-acylation enzyme in *B. theta* and, as such, we have annotated it as lipoprotein *<u>N</u>*-acyltransferase in *<u>B</u>acteroides* (Lnb).

**Figure 2.**
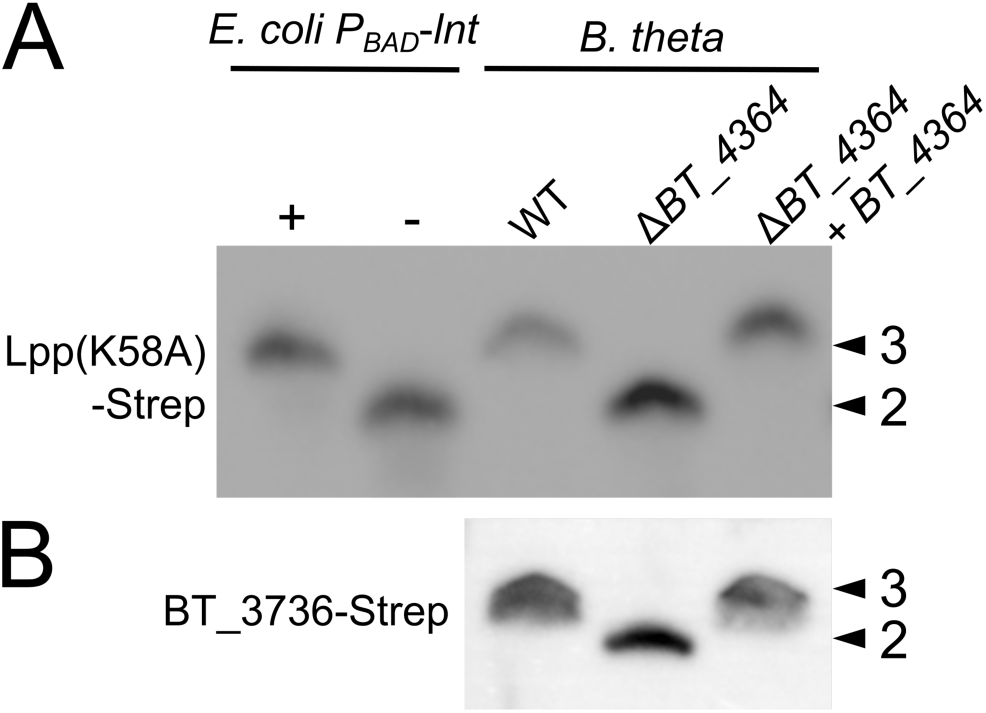
Evidence of lipoprotein *N*-acyltransferase activity by BT_4364 in *B. theta*. Immunoblots against Lpp(K58A)-Strep (A) and BT_3736-Strep (B) using whole-cell lysates of *B. theta* WT, Δ*BT_4364*, and the complemented strain. Lysates from *E. coli* KA349 (*P_BAD_-lnt*) grown under inducing (+) and non-inducing (-) conditions were included as triacyl and diacyl controls for Lpp(K58A)-Strep. Lipoproteins having three acyl chains (arrowhead marked “3”) and two acyl chains (arrowheads marked “2”) are indicated.

### Lnb shares structural homology with NlpC/P60 superfamily enzymes

A BLAST search of Lnb identifies orthologs (>55% amino acid identity) in 487 Bacteroides/Parabacteroides species, and orthologs (>33% amino acid identity) in 103 reference genomes in RefSeq (36, 37). Despite performing the same *N*-acylation activity, Lnb differs considerably from the well-studied Lnt of Gram-negative *E. coli* and *Pseudomonas aeruginosa*. The canonical Lnt protein (512 amino acids in *E. coli*) contains eight transmembrane helices and a large carbon-nitrogen (CN) hydrolase domain located in the periplasm (Fig. 3A, right) (38–40). At 398 amino acids, Lnb is predicted to have five transmembrane helices and a periplasmic domain containing a DUF4105 (Fig. 3A, left). A DALI search of the Lnb periplasmic domain shares limited homology to TseH (PDB 6v98, Z-score=11.6, 12% identity) (Fig. 3B), an NlpC/P60-family cysteine protease, part of a type VI secretion system in *Vibrio cholerae* (41). Interestingly, one of the two proteins required for lipoprotein *N*-acylation in *S. aureus*, LnsA, is also an NlpC/P60 superfamily enzyme (29). While *S. aureus* LnsA and the periplasmic domain of Lnb have no sequence homology, alignment of their AlphaFold-predicted structures shows significant similarities (Fig. 3B).

**Figure 3.**
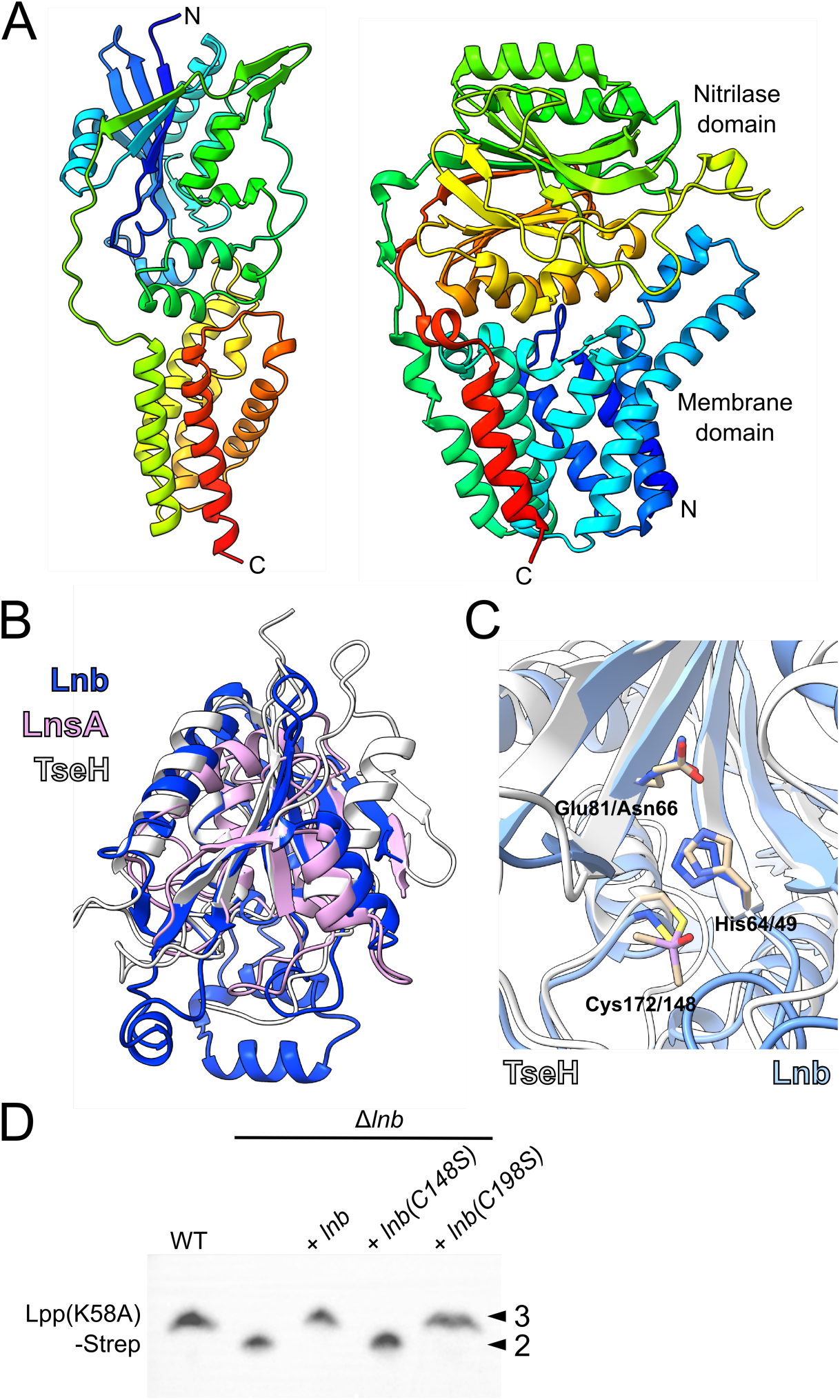
Structural comparisons and active site residue of Lnb. (A) Side-by-side comparison of the AlphaFold-predicted structure of Lnb (left) to the solved structure of *E. coli* Lnt (PDB 5xhq; right), colored rainbow from blue (N-terminus) to red (C-terminus). (B) An overlay of the AlphaFold-predicted periplasmic domain of Lnb (blue), with the AlphaFold-predicted structure of *S. aureus* LnsA (purple) and the solved structure of *V. cholerae* TseH (PDB 6v98; white). (C) Close-up of the active site of TseH (white) overlaid with Lnb (blue). (D) Whole cell lysates of indicated *B. theta* strains were separated by SDS-PAGE and immunoblotted against Lpp(K58A)-Strep. Lipoprotein having three acyl chains (arrowhead marked “3”) and two acyl chains (arrowheads marked “2”) are indicated.

Based on the overlay with the catalytic triad Glu81-His64-Cys172 of TseH (Fig. 3C), we predicted that C148 or C198 could be a catalytic residue of Lnb. We mutated both residues to serine and tested the resulting mutants’ ability to *N*-acylate the reporter Lpp(K58A)-Strep in *B. theta*. Only Lnb(C198S) could restore *N*-acylation activity to Δ*lnb* cells, suggesting that C148 is essential for activity of Lnb (Fig. 3D). Notably, *E. coli* Lnt functions by forming a thioester intermediate between its catalytic cysteine (C387) and acyl chain substrate (42). A multisequence alignment of Lnb homologs from 30 common type strains of various gut Bacteroidota revealed that the putative catalytic triad Asn66-His49-Cys148 from the Lnb model overlay with TseH is completely conserved (Fig. S2).

### Cells lacking Lnb are less viable, exhibit altered morphology, and are outcompeted by WT cells

Lipoproteins are involved in many important cellular processes, including cell envelope integrity and division. As such, deletion of lipoprotein modifying enzymes from Gram-negative bacteria is usually lethal or can result in altered cell morphology (26, 43, 44). Thus, we asked if the loss of lipoprotein triacylation affects cell viability or morphology of *B. theta*. To assess cell viability, WT, Δ*lnb*, and complemented strains were grown in rich media (tryptone-yeast extract glucose, TYG) for 7 and 24 hr, representing exponential and late stationary growth phases, respectively; then OD-normalized cells were plated on solid media. All three strains yielded comparable colony forming units (CFU) from the 24 hr cultures, however less CFU were counted from the 7 hr culture of Δ*lnb* cells than WT and complemented (Fig. 4A), suggesting reduced viability during exponential growth. To assess cell morphology, WT, Δ*lnb*, and complemented strains grown in rich media for 7 and 24 hr were visualized by confocal microscopy. Δ*lnb* cells showed increased filamentation and are on average approximately 64% longer than WT and complemented cells (Fig. 4B and C). Taken together, these data suggest that Δ*lnb* cells are less fit than WT cells. Indeed, when co-cultured 1:1 with WT cells, Δ*lnb* cells significantly decreased in percentage by day 1 and are near undetectable by day 2 (Fig. 4D).

**Figure 4.**
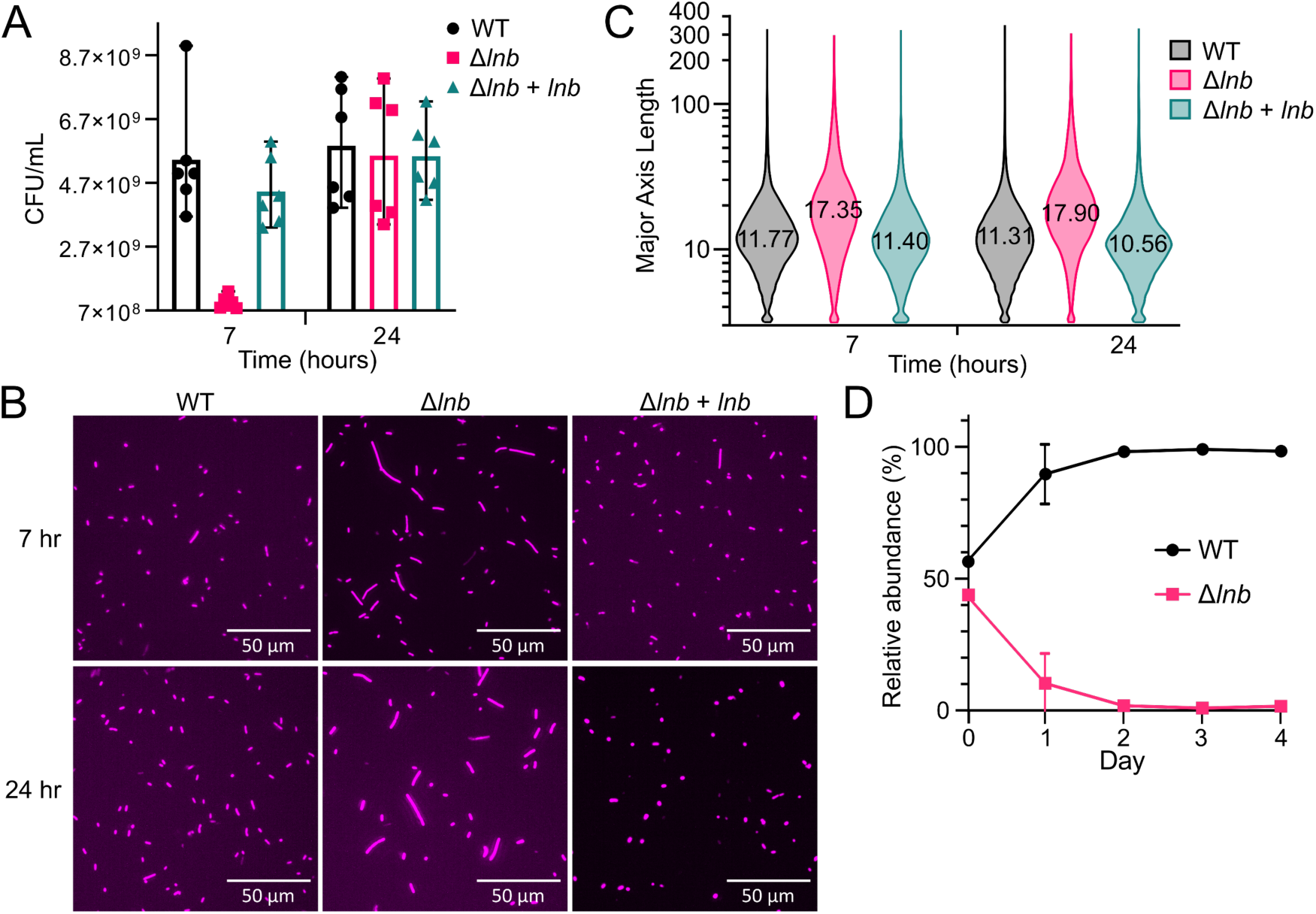
Viability, morphology, and competition of WT versus Δ*lnb* cells. (A) The CFU/mL of WT, Δ*lnb*, and complemented strains when grown in rich media for 7 and 24 hours. Results are from two biological replicates with three technical replicates each. (B) Representative images of WT, Δ*lnb*, and complemented cells grown in rich media for 7 and 24 hours. Cells were stained with CellMask Deep Red, with images taken at 40x magnification. (C) Violin plot quantifying the major axis length of WT, Δ*lnb*, and complemented strains grown in rich media for 7 and 24 hours (*n* = 36,692 to 81,618 objects). The average length is indicated. (D) *In vitro* competitions of barcoded *B. theta* WT and Δ*lnb* strains in rich media. Relative abundance was calculated as the percent composition of a strain’s DNA relative to the total DNA in the sample. Data shown is representative from two biological replicates with three technical replicates each.

### Cells lacking Lnb exhibit growth defects that vary by media composition and carbon source

*Bacteroides* spp. employ distinct lipoproteins involved in uptake, degradation, and import of polysaccharides as carbon sources (2). To examine how loss of lipoprotein triacylation affects growth of *B. theta*, we measured the growth of our strains in rich (tryptone-yeast extract; TY) and minimal media (MM) supplemented with polysaccharides and their cognate monosaccharides. When grown in TY-glucose, a monosaccharide whose uptake and utilization does not rely on lipoproteins, the Δ*lnb* strain grows similar to WT (Fig. 5A). When grown in MM-glucose, however, the Δ*lnb* strain exhibits a lag in growth and does not achieve the same final optical density (OD) as WT (Fig. 5B). Of the monosaccharides we tested, the same trend was observed for fructose, arabinose, and mannose, though not for galactose (Fig. S3A-D). This phenotype is not due to a general inability of the Δ*lnb* strain to grow on monosaccharides, as Δ*lnb* cells grow akin to WT in TY plus monosaccharide (Fig. S3E-H). Taken together, these results suggest that absence of Lnb negatively impacts growth in MM. It is likely that general physiological defects attributed to the loss of lipoprotein *N*-acylation may be exacerbated by growth in MM.

**Figure 5.**
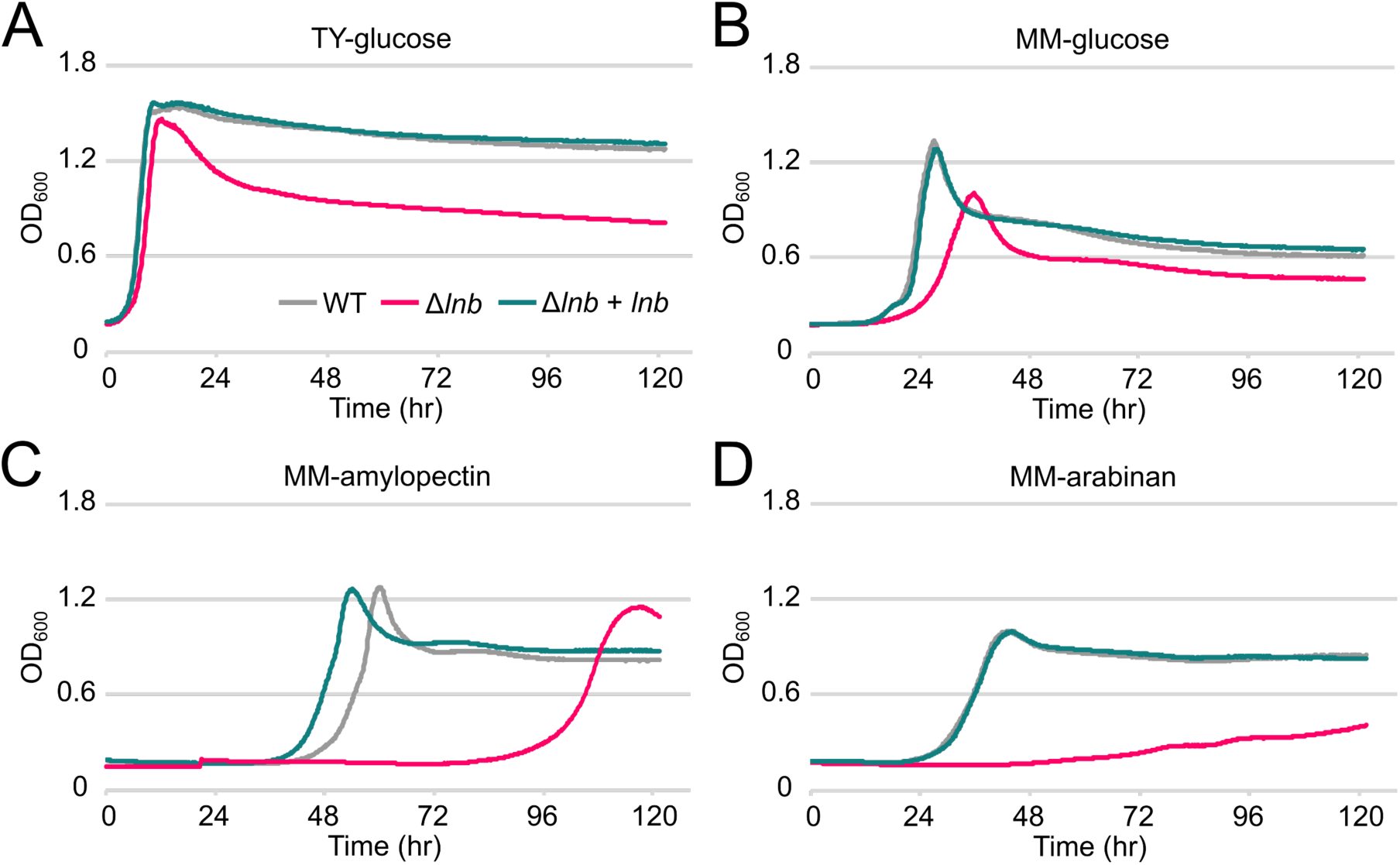
Growth of *B. theta* strains in rich or minimal media with various carbon sources. The optical density at 600 nm (OD_600_) of *B. theta* WT, Δ*lnb*, and complemented strains on TY-glucose (A), MM-glucose (B), MM-amylopectin (C), and MM-arabinan (D) over time. Curves are the average of three technical replicates. An outlier well for Δ*lnb* cells grown in MM-amylopectin was omitted from the data set. The graphs shown are representative of three biological replicates.

In testing growth on more complex polysaccharides in MM, we observed two distinct growth phenotypes of the Δ*lnb* strain that appear dependent on the provided polysaccharide. The first phenotype consists of an initial lag in growth, but similar kinetics to that of WT once the Δ*lnb* strain begins growing. Of the polysaccharides we tested, this was seen with amylopectin (Fig. 4C) and levan (Fig. S4A). The lag was not observed with TY-levan, though a small lag persisted in TY-amylopectin (Fig. S4BC).

The second phenotype is characterized by altered growth kinetics to that of WT, which was observed for Δ*lnb* cells grown in MM-arabinan (Fig. 5D), yeast α-mannan, and mucin *O*-glycans (MOG) (Fig. S4D and S4E). Interestingly, the defect of Δ*lnb* cells was not rescued by growing in TY with the same polysaccharides (Fig. S4F-H).

### Deletion of Lnb reduces cell surface localization of Sus lipoproteins

Canonically, triacylation is prerequisite to lipoprotein OM localization performed by the Lol system (45–47). Thus, we hypothesized that the growth defects observed when Lnb is deleted may be due to mislocalization of diacylated lipoproteins. To investigate this, we immunostained intact WT, Δ*lnb*, and complemented cells for SusE and SusG, two lipoproteins that normally localize to the cell surface during growth on maltosides or starch (5). When fluorescent intensity was quantified, we observed a significant decrease in signal intensity for both lipoproteins on the surface of Δ*lnb* cells compared to WT (an average decrease of approximately 66% for SusE and 38% for SusG; Fig. 6 and Fig. S5A). Δ*susE* and Δ*susG* cells were also imaged as negative controls (Fig. S5B). A Western blot against normalized whole cell lysates confirms that this result is not due to lower expression or protein abundance of SusG (Fig. S6). We cannot conclude from these data where mislocalized SusE or SusG are (i.e. retained in the inner membrane or facing the periplasm from the inner leaflet of the outer membrane) or what step of localization is being perturbed by diacylation (i.e. recognition by LolCDE, surface exposure, etc.).

**Figure 6.**
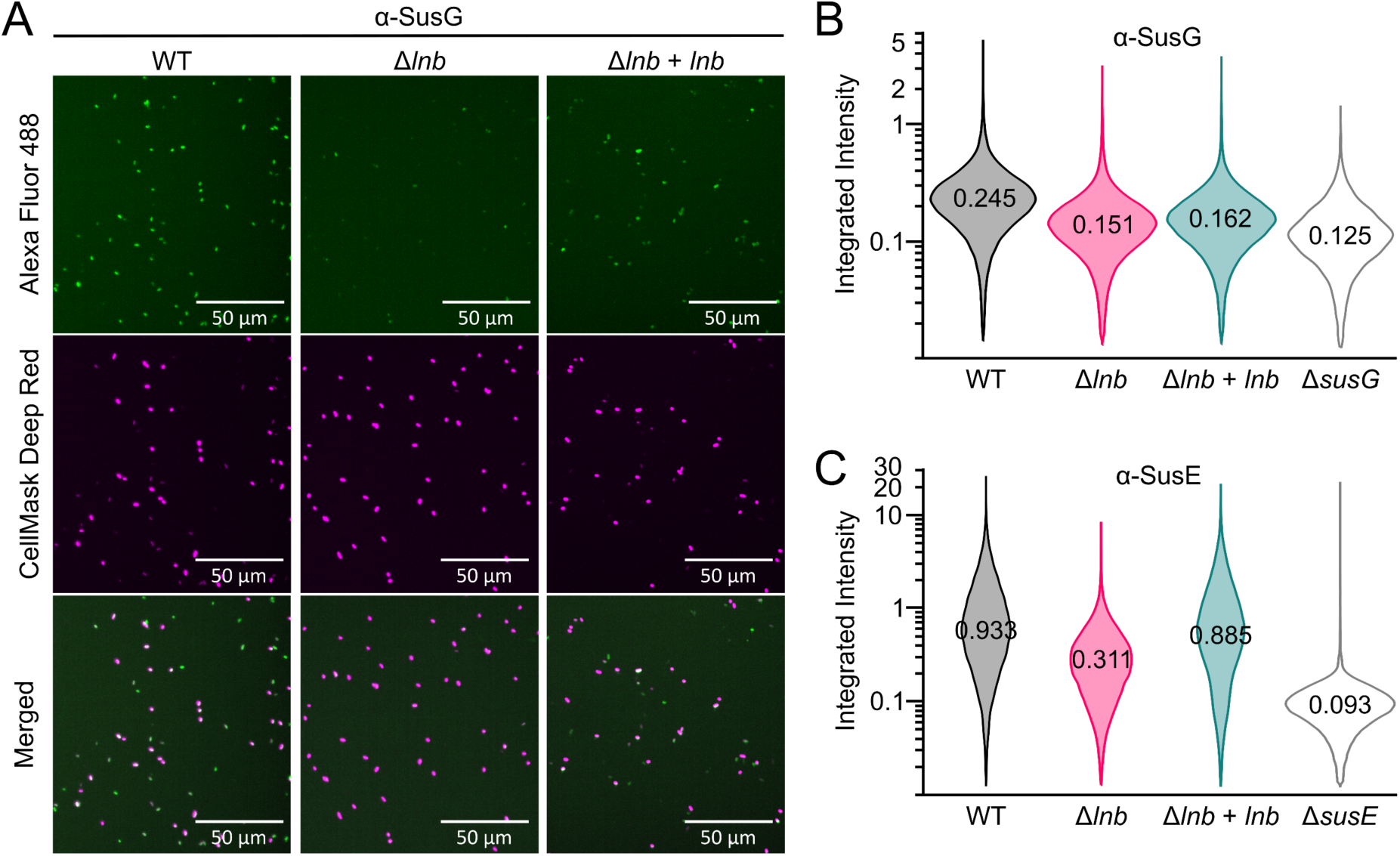
Surface localization of Sus lipoproteins in *B. theta*. (A) Representative microscopy images of *B. theta* WT, Δ*lnb*, and complemented cells when immunostained for SusG. Alexa Fluor 488 and CellMask Deep Red images were taken in the green and red channels, respectively, with equal contrast across all images within the same channel. Quantification of the integrated fluorescent intensity is graphed as a violin plot for SusG (B) and SusE (C). Averages are indicated.

### Deletion of Lnb globally alters membrane and OMV protein composition

Considering our immunostaining data (Fig. 6), we sought a higher throughput and unbiased measurement of lipoprotein localization. To begin, we grew *B. theta* WT and Δ*lnb* strains in MM supplemented with starch, then isolated both membranes and OMVs from each. We chose to include OMVs in our analysis because they are highly decorated with polysaccharide-degrading lipoproteins and can influence cell growth (11, 12). SDS-PAGE analysis of the fractions shows that while the total membrane (TM) protein profiles appear similar between the two strains, the OMV protein profiles from Δ*lnb* differed from that of WT (Fig. 7A).

**Figure 7.**
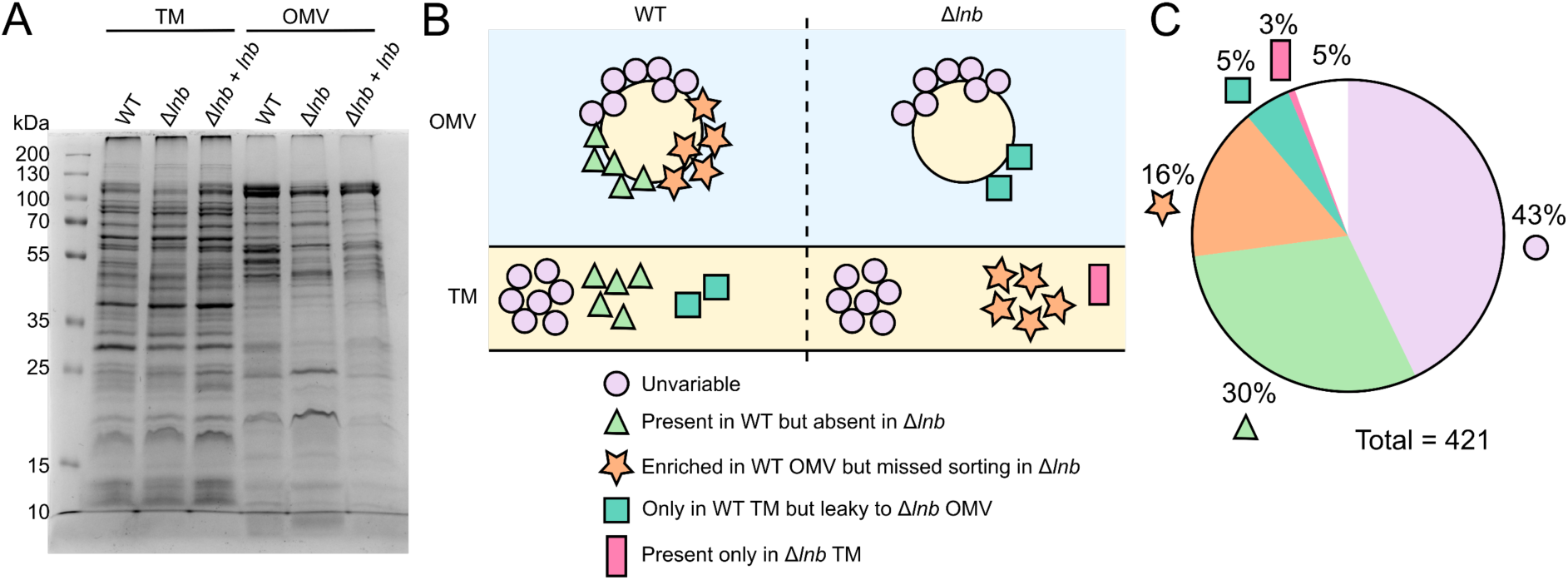
Lnb mutant shows altered TM and OMV protein profiling in *B. theta*. (A) Coomassie blue staining after SDS-PAGE of *B. theta* WT and Δ*lnb* total membrane (TM) and vesicle (OMV) fractions. Equivalent amounts of protein were loaded in each lane (10 μg). (B) Cartoon representation of lipoprotein comportment in *B. theta* WT (left) versus Δ*lnb* (right) in TM and OMVs. Each shape represents approximately 3% of the total lipoproteins detected. (C) Parts-of-a-whole diagram of the same data, following the same legend as in panel B. The white wedge represents unclassified lipoproteins.

To further analyze membrane and vesicle composition, we performed comparative proteomic analyses of these samples (Table S1). Principal component analysis (PCA) showed that the four proteomes clustered separately (Fig. S7), revealing major differences between the groups. Functional analysis of proteins enriched in the TM of WT versus those of Δ*lnb* showed decreased number of proteins associated with energy production and conversion, amino acids and coenzyme transport and metabolism, as well as translation, ribosomal structure and biogenesis (Table S2). These results suggest a global decrease in metabolism when Lnb is deleted and may partially account for the growth defects observed even when grown in rich media (Fig. S4).

In analyzing lipoproteins specifically, we observed a significant reduction in the quantity of lipoproteins detected in Δ*lnb* cells compared to WT. Of the 421 total lipoproteins identified in the proteomics, approximately 30% (124/421) were absent from Δ*lnb* cells (Fig. 7B and C; full dataset in Table S1).

Differences in lipoprotein sorting between TM and OMVs were also observed between WT and mutant. Of 86 lipoproteins showing altered sorting, 20 lipoproteins were retained in the TM of WT but were “leaky” to the OMVs of Δ*lnb* (5%), while 66 lipoproteins enriched in the OMVs from WT (16%) were not enriched in OMVs from the mutant. Surprisingly, a minor portion of lipoproteins was present only in Δ*lnb* (3%, 11/421). Despite these differences, a large portion of lipoproteins were unaltered between the two strains (42%, 177/421). As analysis was performed on total membranes, these data do not indicate protein localization in the inner versus outer membrane. It is currently unclear what factor(s) determine which lipoproteins are mislocalized when Lnb is deleted.

## Discussion

Lipoproteins are ubiquitous components of bacterial cell envelopes and carry out an array of functions essential to survival. Bacteroidota in particular deploy cell-surface lipoproteins that capture and degrade various polysaccharides, contributing to their persistence in the human gut and influence on host health and disease. Canonically in Gram-negative bacteria, lipoproteins are sequentially modified by the diacylglycerol transferase Lgt and signal peptidase II Lsp, then triacylated by the *N*-acyltransferase Lnt. Overall, however, empirical validation of lipoprotein acylation and biosynthetic pathways across bacteria is lacking. Indeed, studies on various Gram-positive species revealed more lipoprotein forms and lipoprotein-modifying enzymes than previously thought existed (29–31). While empirical evidence supports *N*-acylation of lipoproteins in *B. fragilis* (14), no prior study described how *N*-acylation occurs in this species. Herein, we show that lipoproteins in *Bacteroides* spp. are not *N*-acylated by Lnt, but rather by a previously undescribed protein that we named Lnb.

Lipoprotein *N*-acylation is widely regarded as essential in Gram-negative bacteria, being crucial to proper recognition by the LolCDE exporter and subsequent localization to the OM (21). Nevertheless, we were able to delete Lnb from *B. theta*, resulting in diacylated lipoproteins (Fig. 2). In *E. coli*, lethality resulting from the loss of Lnt is largely attributed to mislocalization of diacylated Lpp to the inner membrane (22). Deletion of Lpp or mutation to its peptidoglycan-crosslinking amino acid (K58) can partially restore growth when Lnt is depleted in these genetic backgrounds (48, 49). *Bacteroides* spp., however, do not encode an identifiable ortholog to Lpp (50), possibly allowing for deletion of Lnb.

Moreover, Lnt can be successfully deleted in some Gram-negative species that encode an alternate Lol system component (LolF) that exports diacylated lipoproteins to the OM (24, 26, 51), though it is currently unclear if *Bacteroides* encodes LolF.

While our immunostaining (Fig. 6) and proteomics (Fig. 7) show differences between WT and Δ*lnb* cells, overall some portion of lipoproteins are still localized to the cell surface when Lnb is absent. This suggests that lipoprotein *N*-acylation is not solely for proper OM localization, as further evidenced by the existence of *N*-acylated lipoproteins among Gram-positive bacteria lacking an OM, such as some *Staphylococcus* spp. (28). Instead, lipoprotein *N*-modification has been proposed to be a “post-edit” to the conserved Lgt-Lsp pathway, with different bacterial species evolving their own methods for *N*-modification (29, 30, 52). That Lnb appears phylogenetically confined to *Bacteroides* spp. supports this theory. The selective pressure(s) driving lipoprotein *N*-modification are uncertain, though a role in resistance to copper may be one factor (52, 53). Structural similarities between Lnb and LnsA of *Staphylococcus* spp. (Fig. 3B), both NlpC/P60 superfamily enzymes, support the establishment of a new family of lipoprotein *N*-acyltransferases and warrant further research into other species encoding such genes.

Although Lnb can be deleted from *B. theta*, its deletion negatively impacts cell physiology. Δ*lnb* cells are less viable and are outcompeted by their WT counterparts and display altered cell morphology (Fig. 4). It is likely that there are other phenotypes resulting from the deletion of Lnb that we did not explore in this study, such as capsule and peptidoglycan formation, and outer membrane stability and permeability. Notably, Δ*lnb* cells grow relatively well on some polysaccharides but not on others, even in rich media (Fig. 5, Fig. S3 and S4). The complexity of the polysaccharide and the suite of lipoproteins involved in its utilization are likely factors to the observed growth phenotypes. As a portion of lipoproteins indeed reach the cell surface in Δ*lnb* cells (Fig. 6), cell growth may be slow until enough polysaccharide is degraded to allow more robust growth. It is also possible that Δ*lnb* cells are more prone to lysis due to membrane defects, in turn releasing polysaccharide-degrading proteins that feed intact cells. More studies are needed to explore these hypotheses.

Relatedly, altered OMV composition likely also influences growth. When grown in MM with starch, Δ*lnb* cells produce OMVs, however their lipoprotein cargo differs significantly from those produced by WT (Fig. 7). Similar results are expected for Δ*lnb* cells grown in rich media or MM supplemented with other carbon sources, though the individual proteins identified would vary, as observed for *B. theta* WT by Sartorio et al. (12). In general, an astounding 30% of lipoproteins were absent from Δ*lnb* cells, another 20% showed altered sorting between TM and OMV, and yet still nearly half of lipoproteins were unchanged between WT and Δ*lnb*. These results may indicate that *N*-acylation is required for certain lipoproteins but is less important for others. An alternate pathway(s) for lipoprotein transport is also possible, as has been suggested in *E. coli* (54). Nevertheless, our data indicate widespread alterations in lipoprotein abundance and sorting, including proteins beyond those necessary for polysaccharide utilization, highlighting the fundamental importance of Lnb.

Lipoprotein biosynthesis has been suggested as a target for novel antibiotics (55–57). However, some bacteria tolerate losing their lipoprotein biosynthetic enzymes, especially amongst Gram-positive organisms (55, 58, 59). Indeed, this is also true for *B. theta*, albeit with negative outcomes for the cell, notably the inability to grow on certain polysaccharides such as MOG (Fig. 4, Fig. S4). *Bacteroides* is one of the main genera implicated in intestinal mucus degradation that can lead to colitis (60, 61). Thus, development of an antibiotic against Lnb would target mucin-degrading *Bacteroides* spp. in a highly specific manner, since Lnb is phylogenetically confined to *Bacteroides*. Further studies are required to characterize Lnb at the enzymatic level and within other *Bacteroides* strains.

## Methods

### Bacterial strains and growth conditions

The strains in this study are listed in Table S3. *E. coli* strains were derived from reference strain BW25113 and grown aerobically in LB medium at 37°C with agitation. Cultures were supplemented with 0.2% (wt/vol) L-arabinose or 0.2% (wt/vol) D-glucose where indicated. Antibiotic markers were selected with carbenicillin (100 µg/mL), kanamycin (25 µg/mL), and spectinomycin (50 µg/mL).

*B. theta* strains were derived from VPI-5482 Δ*tdk* (herein referred to as wildtype) and were routinely cultured in a Coy anaerobic chamber at 37°C with an atmosphere of 10% H_2_, 5% CO_2_, 85% N_2_. Strains were grown in either Brain heart infusion (BHI) medium supplemented with 5 μg/ml hemin and 1 μg/ml vitamin K_3_, or tryptone-yeast extract-glucose (TYG) medium (62). When applicable, strains were grown in *Bacteroides* minimal medium (MM) containing 100 mM KH_2_PO_4_ (pH 7.2), 15 mM NaCl, 8.5 mM (NH_4_)_2_SO_4_, 4 mM L-cysteine, 1.9 mM hematin/200 mM L-histidine (prepared together as a 1,000x solution), 100 mM MgCl_2_, 1.4 mM FeSO_4_.7H_2_O, 50 mM CaCl_2_, 1 μg/mL vitamin K_3_ and 5 ng/mL vitamin B_12_. Carbohydrates were supplemented to a concentration of 5 mg/mL, except MOG (10 mg/mL). When applicable, gentamicin was used at 200 μg/mL, erythromycin at 25 μg/mL, and tetracycline at 3 μg/mL.

For growth experiments in a plate reader, overnight cultures were washed thrice in phosphate buffered saline (PBS) and normalized to a starting OD_600_ of 0.05 in MM plus the experimental carbohydrate.

Growth experiments were performed in triplicate at 37°C in 96-well plates and the OD_600_ was recorded every 10 minutes. The averages are reported in each figure. Three biological replicates were performed.

### Construction of plasmids, *E. coli* Δ*lnt* strain, and *Bacteroides* strains

All plasmids (except the genomic library, described below) were constructed via FastCloning (63) or with the In-Fusion HD Cloning Kit (Takara). Primers used in this study are listed in Table S4. The *lnt::specR* allele was transduced from strain TXM1036 (30)(a gift from Dr. Timothy Meredith) into recipient strains using P1*vir* (a gift from Dr. Lydia Freddolino) when appropriate using standard protocols (64). Deletions in *B. theta* were created using the Δ*tdk* allelic exchange method (65). Overexpression of Lpp(K58A)-Strep and BT_3736-Strep were driven by the strong promoter of pWW1376, a gift from Dr. Justin Sonnenburg (35), integrated at an *att* site. For complementation of BT_4364 to the deletion strain, the constitutive promoter *us1311* was engineered into pWW1376 for dual or single expression.

### Construction of *B. fragilis* genomic library and complementation method

Following the protocol of Cho 2019 (66), genomic DNA from *Bacteroides fragilis* NCTC 9343 was digested with *Sau*3AI at 20°C for an optimized time of 40 min. After separation by gel electrophoresis, DNA fragments ranging from 3 to 5 kb were isolated. Fragments were then ligated with T4 ligase into pUC19 vector that had been linearized with *Bam*HI and dephosphorylated with shrimp alkaline phosphatase (rSAP; NEB). The resulting plasmids were transformed into Stellar Competent *E. coli* cells (Takara Bio) and plated on LB agar plus carbenicillin. Transformants were scraped from the agar surface into 1 mL PBS, with the final library then isolated from 100 µL of these cells.

The library was transformed into the conditionally lethal *E. coli lnt* mutant KA349 (30) and plated on LB agar containing carbenicillin and glucose. A portion of the transformed cells was also plated on LB agar containing arabinose as a positive control. Transformants that grew in the absence of arabinose were substruck to confirm phenotypic rescue. Plasmids isolated from the positive colonies were sequenced to identify the *B. fragilis* gDNA insert.

### Colony formation by Lnt-depleted KA349 transformed with experimental plasmids

KA349 cells grown in LB plus kanamycin and arabinose were washed thrice in PBS then seeded 1:1000 into fresh LB plus kanamycin and glucose. At OD_600_ ∼0.5-0.6, cells were made competent and transformed via heat-shock with 50 ng of plasmid. After 1 hr outgrowth with no arabinose or glucose, 100 µL each was spread onto three LB agar plates containing arabinose and three containing glucose. Plates also contained carbenicillin for transformant selection and 10 µg/mL vancomycin to reduce background growth (30).

CFU were enumerated the following day. Three biological replicates were performed.

### Immunoblotting

Whole cell lysates were separated by SDS-PAGE with a 16.5% Tris-tricine gel (67) for Lpp(K58A)-Strep and BT_3736-Strep, and a 10% Tris-glycine gel for SusG. Proteins were transferred to polyvinylidene difluoride (PVDF) membrane (0.2 µm). For Lpp(K58A)-Strep and BT_3736-Strep, the membrane was incubated with a 1:5,000 dilution of StrepTactin-horseradish peroxidase (HRP) conjugate (Bio-Rad), and 1:500 dilution of custom polyclonal antibodies (Cocalico Biologicals) for SusG. Goat anti-rabbit IgG HRP conjugate (Bio-Rad) was used as a secondary at 1:5,000 dilution when necessary. Signals were detected by enhanced chemiluminescence.

### Affinity purification of tagged lipoproteins

Lpp(K58A)-Strep was purified from *E. coli* as previously described (30), with cells lysed by sonication.

### MALDI-TOF MS and MS/MS

Preparation of lipoproteins for N-terminal structural characterization by MALDI-TOF MS was described previously (68). Briefly, purified Lpp(K58A)-Strep was separated by SDS-PAGE (15% Tris-glycine), transferred to nitrocellulose membrane, then stained with Ponceau S. Bands were excised, washed with water, and proteolyzed with trypsin at 37°C overnight. The nitrocellulose was then sequentially washed with 0.1% trifluoroacetic acid (TFA), 10% acetonitrile, and 20% acetonitrile. A final incubation with 10 mg/mL α-cyano-4-hydroxycinnamic acid (CHCA) in 2:1 chloroform-methanol eluted the final tryptic N-terminal lipopeptides. MS spectra were collected on an UltrafleXtreme MALDI-TOF (Bruker Daltonics) mass spectrometer in positive reflectron mode. MS/MS spectra were acquired on the same instrument in LIFT mode.

### Viable cell counting

Overnight cultures of *B. theta* were inoculated into fresh TYG to an OD_600_ of 0.05. After 7 and 24 hr of growth, 1 mL of cells were removed and the OD_600_ normalized to 1.9. Cells were serial diluted and the 10^-5^, 10^-6^, and 10^-7^ dilutions were used to quantify CFU via drip plating method. Briefly, 20 µL of each dilution was spotted onto TYG-agar then the plate rotated 90° vertically to allow spots to run down the agar surface. Colonies were counted after ∼36 hr incubation and the CFU/mL calculated based on the dilution factor. Two biological replicates with three technical replicates each were performed.

### Cell competition and qPCR

pNBU2-based plasmids encoding isogenic barcode 1 or 14 (gifts from Dr. Eric C. Martens)(69) were integrated into *B. theta* WT and the Δ*lnb* mutant, respectively. Strains were grown overnight in TYG, normalized to an OD_600_ of 0.5, then mixed in a 1:1 ratio. One mL of the mix was reserved while the rest was diluted to an OD_600_ of 0.05 into fresh TYG and divided into replicates of 2 mL each. At 24 hr intervals for 5 days, cells were passaged 1:100 into fresh TYG, with an aliquot from each passage collected and frozen. Genomic DNA was isolated from these aliquots using the Qiagen DNeasy Blood and Tissue kit, quantified via Nanodrop (Thermo Scientific), and normalized to 10 ng/µL. Quantitative PCR (qPCR) was carried out in a CFX Connect Real-Time System (Bio-Rad) using a homemade SYBR mastermix for 40 cycles of 95°C for 3 s, 55°C for 20 s, 72°C for 20 s, followed by a melting step to determine amplicon purity. Forward primers were specific to barcodes 1 or 14 and were paired with a universal reverse primer. Samples were normalized to a DNA standard curve of genomic DNA from each respective strain. Relative abundance was calculated as the percent composition of a strain’s DNA relative to the total DNA in the sample.

### Preparation of cells for microscopy

Overnight cultures of *B. theta* WT, Δ*lnb*, and complemented cells were passaged into fresh TYG to an OD_600_ of 0.05. Aliquots were removed at 7 and 24 hrs, washed twice with PBS, and resuspended in 250 µL PBS. Cells were then incubated for 1.5 hr at room temperature with 750 µL of 6% formalin. Cells were again washed twice in PBS, then resuspended in 50 µL of a 1:5,000 dilution of CellMask Deep Red (Thermo Fisher Scientific) and incubated for 30 min in the dark. Cells were washed thrice in PBS and resuspended to 1 mL. Three µL of a 1:100 dilution of the cells were deposited onto slides for imaging.

### Immunostaining

Overnight cultures of *B. theta* WT, Δ*lnb*, and complemented cells were diluted into MM with maltose to an OD_600_ of 0.05. Cells were removed once they reached an OD_600_ of 0.7, washed twice with PBS, fixed in formalin as above, then washed twice again. Cells were resuspended in 500 µL blocking solution (2% goat serum and 0.02% NaN_3_ in PBS) and rocked overnight at 4°C. After washing, cells were incubated with a 1:500 dilution of anti-SusG and 1:5,000 dilution of anti-SusE antibodies in blocking solution for 2 hr at room temperature. Following two washes in PBS for 10 min each, cells were rocked in 0.4 mL of a 1:500 dilution of goat anti-rabbit IgG Alexa Fluor 488 secondary antibody (Invitrogen) in blocking solution for 1 hr in the dark. Cells were again washed in PBS for 10 min each, stained in 50 µL of CellMask Deep Red as above, and washed a final three times before deposition onto slides for imaging.

### Image acquisition and analysis

Cells were imaged using a CellVoyager CQ1 Benchtop High-Content Analysis System (Yokogawa) with a 40x lens. Alexa Fluor 488 and CellMask Deep Red images were taken in the green and red channels with exciting wavelengths of 488 nm and 659 nm, respectively, with spinning disk confocal and 500 ms exposure times (70). Laser power was adjusted to yield an optimal signal-to-noise ratio for each channel. Laser autofocus was used and 100 fields per slide were imaged. Maximum intensity projection images were collected from 25 confocal planes with a 2.5 µm step size. Bacterial cells were segmented using Otsu two-class thresholding with CellProfiler using the CellMask Deep Red channel and intensity and size/shape features were tabulated for the Alexa Fluor 488 and CellMask Deep Red channels (71). Cell-level data were preprocessed and analyzed in the open source Knime analytics platform. Data analysis was completed using Microsoft Excel and GraphPad Prism. A paired student t-test was performed to determine the significant differences in Alexa Fluor 488 intensity for each strain.

### Subcellular fractionation

OMVs were purified by ultracentrifugation of filtered spent media. Briefly, 50 mL of *B. theta* cultures from early stationary phase were centrifuged at 6,500 rpm at 4°C for 10 min.

Supernatants were filtered using a 0.22-μm-pore membrane (Millipore) to remove residual cells. The filtrate was ultracentrifuged at 200,000 x*g* for 2 hr (Optima L-100 XP ultracentrifuge; Beckman Coulter). Supernatants were discarded, and the pellet resuspended in 50 mM HEPES. Purified OMV preparations were lyophilized for MS analysis.

Total membrane preparations were performed by cell lysis and ultracentrifugation. Cultures from early stationary phase were harvested by centrifugation at 6,500 rpm at 4°C for 10 minutes. The pellets were gently resuspended in 50 mM HEPES containing complete EDTA-free protease inhibitor mixture (Roche Applied Science) followed by cell disruption. Centrifugation at 6,500 rpm at 4°C for 5 min was performed to remove unbroken cells. Total membranes were collected from the resulting supernatant by ultracentrifugation at 200,000 x*g* for 1 hr at 4°C. Total membrane fractions were lyophilized for MS analysis. Membrane and vesicles fractions were analyzed by standard 10% Tris-glycine SDS-PAGE. Protein concentrations were determined using a DC Protein Assay kit (Bio-Rad).

### Sample preparation for proteomic analyses

Lyophilized protein samples were solubilized in 4% SDS, 100 mM HEPES by boiling for 10 min at 95°C. Protein concentrations were determined by bicinchoninic acid protein assay (Thermo Fisher Scientific) and 200 μg of each biological replicate prepared for digestion using Micro S-traps (Protifi, USA) according to the manufacturer’s instructions. Briefly, samples were reduced with 10 mM DTT for 10 min at 95°C then alkylated with 40 mM iodoacetamide (IAA) in the dark for 1 hr. Samples were acidified to 1.2% with phosphoric acid and diluted with seven volumes of S-trap wash buffer (90% methanol, 100 mM tetraethylammonium bromide pH 7.1) before loading onto S-traps and washing three times with 400 μL of S-trap wash buffer. Samples were then digested with 2 μg of trypsin (a 1:100 protease/protein ratio) in 100 mM tetraethylammonium bromide overnight at 37°C. The digests were collected by centrifugation then sequentially washed with 100 mM tetraethylammonium bromide, 0.2% formic acid, and 0.2% formic acid/50% acetonitrile. Samples were dried down and further cleaned using C18 Stage (72, 73) tips to ensure the removal of any particulate matter.

### Reverse phase liquid chromatography-mass spectrometry (LC-MS)

Prepared C18-purified peptides from each sample were resuspended in Buffer A* (2% acetonitrile, 0.01% trifluoroacetic acid) and separated using a two-column chromatography setup composed of a PepMap100 C_18_ 20-mm by 75-μm trap and a PepMap C_18_ 500-mm by 75-μm analytical column (Thermo Fisher Scientific). Samples were concentrated onto the trap column at 5 μL/min for 5 min with Buffer A (0.1% formic acid, 2% DMSO) and then infused into an Orbitrap Fusion Lumos equipped with a FAIMS Pro interface at 300 nL/min via the analytical columns using a Dionex Ultimate 3000 UPLCs (Thermo Fisher Scientific). 125-minute analytical runs were undertaken by altering the buffer composition from 2% Buffer B (0.1% formic acid, 77.9% acetonitrile, 2% DMSO) to 22% B over 95 min, then from 22% B to 40% B over 10 min, then from 40% B to 80% B over 5 min. The composition was held at 80% B for 5 min, and then dropped to 2% B over 2 min before being held at 2% B for another 8 min. The Fusion Lumos Mass Spectrometer was operated in a stepped FAIMS data-dependent mode at two different FAIMS CVs -40 and -60. For each FAIMS CV a single Orbitrap MS scan (300-1600 m/z and a resolution of 60k) was acquired every 1.5 sec followed by Orbitrap MS/MS HCD scans of precursors (fixed NCE 35%, maximal injection time of 55 ms and a resolution of 15k).

### Proteomic data analysis

Identification and LFQ analysis were accomplished using MaxQuant (v2.0.2.0) (74) with the *Bacteroides thetaiotaomicron* VPI-5482 proteome (UniProt: UP000001414) allowing for oxidation on methionine. Prior to MaxQuant analysis, data files were separated into individual FAIMS fractions using the FAIMS MzXML Generator (75). The LFQ and “Match Between Run” options were enabled to allow comparison between samples. The resulting data files were processed using Perseus (v1.4.0.6) (76) to compare the growth conditions using student t-tests as well as Pearson correlation analyses. Classification of proteins identified by MS were performed using UniProt (77), PSORT (78), SignalP (79), and PULDB (80). Functional analysis was performed using eggNOG-mapper (PMID: 30418610).

## Supporting information

Supplemental Table S1 Proteomics

## Data availability

The mass spectrometry proteomics data has been deposited in the Proteome Xchange Consortium via the PRIDE partner repository with the data set identifier PXD043219 (reviewer login details: username, password: m9jU1Hf0). All plasmids, proteins, bacterial strains, and other reagents generated for this work will be made freely available to researchers using them for non-commercial reasons.

## Author contributions

K.M.A.: conceptualization, funding, data collection and analysis, project administration, supervision, writing – original draft. J.J.: data collection and analysis, visualization, writing. M.G.S.: data collection and analysis, visualization, writing. N.E.S.: data collection. J.M.P.: data collection and analysis, visualization, writing. J.Z.S.: data analysis and resources. M.F.F.: funding. N.M.K.: conceptualization, funding, project administration, supervision, data analysis, visualization, writing.

## Acknowledgements

This work was supported by NIH grants R01DK125445 (to N.M.K.), and R21AI151873 and R21AI168719 (to M.F.F.). K.M.A. is partially supported by the Michigan Life Sciences Fellows program. N.E.S. is supported by an Australian Research Council Future Fellowship (FT200100270) and an ARC Discovery Project Grant (DP210100362). We thank the Melbourne Mass Spectrometry and Proteomics Facility of The Bio21 Molecular Science and Biotechnology Institute, as well as the Mass Spectrometry and Metabolomics Core at Michigan State University, for access to MS instrumentation.

