## Supplemental information for "Identification and Characterization of the Lipoprotein *N*-acyltransferase in *Bacteroides*"

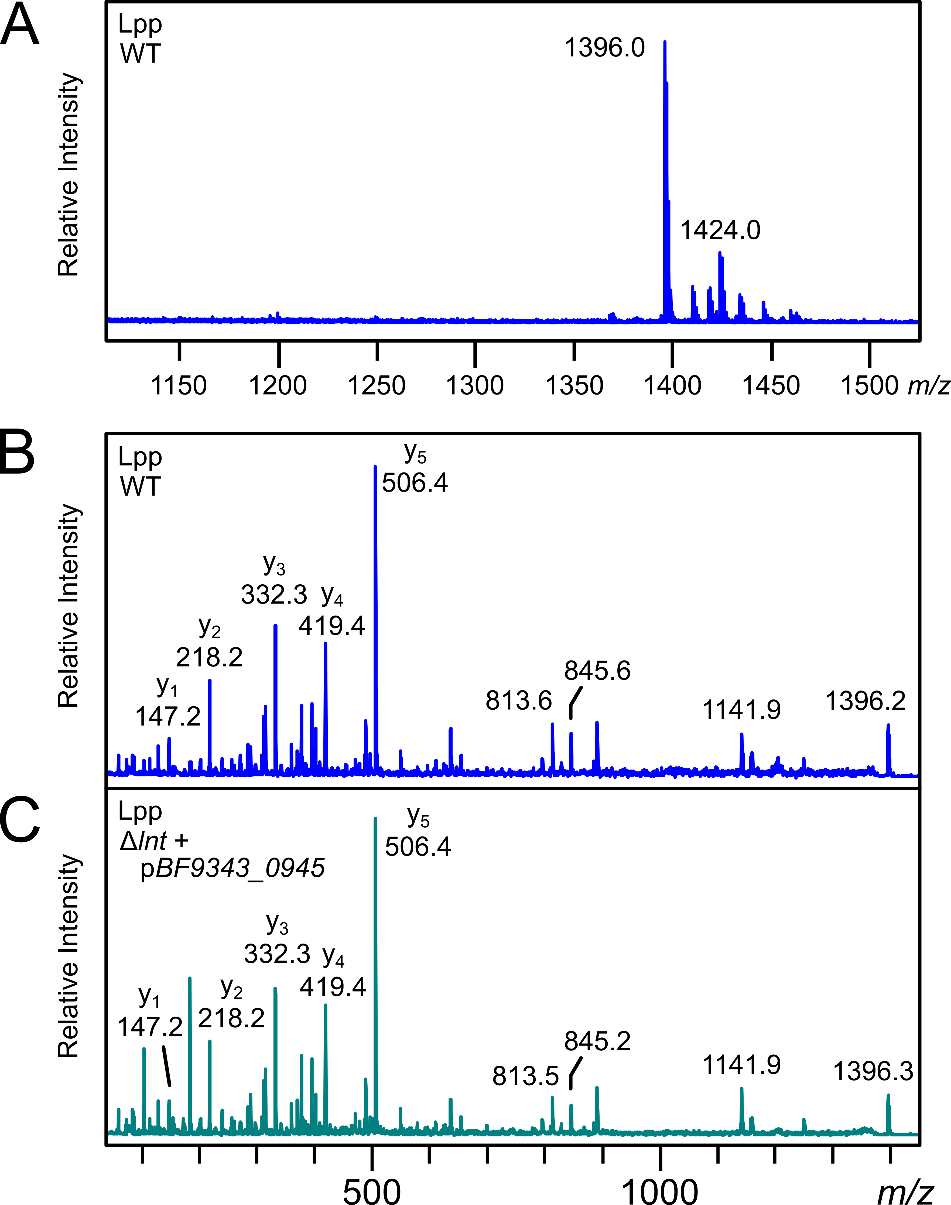

Fig. S1 (A) Trypsinized lipopeptides of Lpp purified from wildtype *lnt E. coli* cells were eluted from nitrocellulose and analyzed by MALDI-TOF MS. The prominent ion of *m/z* 1396.0 is consistent with the triacylated tryptic N-terminus of Lpp. MS/MS analysis of parent peak *m/z* 1396 of Lpp purified from wildtype *lnt E. coli* (B) and Δ*lnt* + p*BF9343_0945* cells (C) show characteristic peaks at *m/z* 813 and 845 corresponding to the *N*-acyl (C16:0)-dehydroalanyl peptide.

A. CLUSTAL O(1.2.4) multiple sequence alignment of Lnb homologs

```
Osplan         MQVLLFLHRLYFFVILNIRNKKTYKMKKIIFLFI--ISS-LIS-----------------	40
Barn_int       -------------------------MKKKLYIILGFILVVLFAN-EKIQAQ---------	25
Pdista         -------------------------MTKSIFLFLLGILSLSAM-----------------	18
Pjohns         -------------------------MERYIYALLLLFASVSLQ-----------------	18
Pmerdae        -------------------------MKKNIFLLFLLFTAVSLQ-----------------	18
Pgold          -------------------------MKKNLLLLFLFLCLIPAK-----------------	18
Pgord          -------------------------MKKYLLLLFLLLLLVPAK-----------------	18
Dgadei         -------------------------MPKLILLLTLII-GFTAQT-K---AQ---------	21
Dmossii        -------------------------MRNLFFIVFLIIASFSQGL-RSQQNA---------	25
Pcopri         -------------------------MKRFKHIFSAILAVILINLTTEVAAQDFSRDKDSD	35
Begger         -------------------------MK---QYLVFLLSLLTCLPCSSTATA---------	23
Bclarus        -------------------------MKFSLPAFRILIFLFLCLPFIAVQAQ---------	26
Bsterc         -----------------------------------------------MQAQ---------	4
Bunif          -------------------------MKRSLLIHIYIVIFFFCLPAANAQV----------	25
Bfluxus        -------------------------MKQNLYISLYLIFFLLCPYPAIAQM----------	25
Boleici        -------------------------MKRTILLLIISAFCAI-------------------	16
Bcell          -------------------------MKRNLLILIYFLLRLLCPSTASAQP----------	25
Bintesti       -------------------------MKRKLLISVYFILCSLCPHIATAQE----------	25
Bfrag          -----------------------MNSKLRH---LLLIVFSIFPILTWG------------	22
Bfineg         -----------------------------M---LFFST-TDVSAMKSN------------	15
Btheta         ------------------------M-KRTI---LYPLLTLYLLIFSNG------------	20
Bcaccae        ------------------------M-KRSF---LYVIFSLFLLLLPIG------------	20
Bxylani        ------------------------M-KRFF---LYTILSFFLLLPSAG------------	20
Bovatus        ------------------------M-KRFF---SYTILSFFLLLLPTG------------	20
Bnordii        -----------------------MM-KKIK---IILILFFLCLATNGK------------	21
Bsalyer        ------------------------M-KRIK---TILILFLVCLTAKGE------------	20
Pplebeius      -------------------------MKKKFVFLLC-ATLISCVFSSTKVWG---------	25
Pmassi         -------------------------MKKLLVLL-Y---L---------------------	10
Pdorei         -------------------------MNKLIIAL-I-LLF---------------------	12
Bvulg          -------------------------MNKLIIAL-I-LLF---------------------	12


Osplan         -------IKGIGLTLSPETEVSILTCAPG-NELYSLFGHTAIRINDP-RHQIDRVYNYGT	91
Barn_int       --------ESKPGILPDTLQVSLLTCGPG-TEVYELFGHTALRVKQQRPGGFDYVFNYGM	76
Pdista         ---------AQ-PKLSEEARISLMTSAPYDEEVFTVYGHAALRIYDP-KQNIDYIFNYGI	67
Pjohns         ---------AQYPLLSKDAEISLLTVSPSEDEVYTVYGHTALRVRDT-SKKLDTVFNYGI	68
Pmerdae        ---------AQYPLLSKDAEISLLTVSPSEDEVYTVYGHTALRVRDA-SKKLDTVFNYGI	68
Pgold          ---------AQ-TGLSDEAQISILTAAPSDDEVYTVYGHTAIRVKDT-LRKLDTVFNYGI	67
Pgord          ---------AQ-IKLSDEAQISILTAAPSDEAVFTLYGHTAVRVKDS-LHKIDLVFNYGI	67
Dgadei         --------EIQRIVLSDSAKVSLLTNAPWDEAVYSLFGHTSMRISDP-TQNIDYAFNFGL	72
Dmossii        --------FRSPITLSDTAQISLLTSSAWEKEIYALFGHTAIRVCDT-TQHLDVVFNYGL	76
Pcopri         ISVQNSRVNEEATAWLDSVDISLLTCGPG-QEVWSYYGHTALRIQNK-AMGTDVAVNWGM	93
Begger         ----HTERAAKASPTADSIRISLLTCASG-EEIYSLFGHTAIRYENY-TRGIDAVFNYGI	77
Bclarus        ---------EHKRDTPDSIRISLLTCASG-EEIYSLFGHTAIRYENH-TRGIDAVFNYGI	75
Bsterc         ---------EPKRVIPDSIHINLLTCASG-EEIYSLFGHTAIRYENY-TRGIDAVFNYGI	53
Bunif          -------TQQQKSAATDSVRVSLLTCAAG-GEIYSLFGHTAIRYENY-TRGIDAVFNYGM	76
Bfluxus        --------QAQRESTPDSIRISLLTCAPG-GEIYYLFGHTAIRYENL-TRGIDAVFNYGI	75
Boleici        ---------GLKAQRADSIRISLLTCASG-GEIYSLFGHTAIRYENY-TQGIDAVFNYGM	65
Bcell          --------QTPQEATPDSIRISLLTCASG-GEIYSLFGHTAIRYENF-TRGIDAVFNYGM	75
Bintesti       --------QTPREKITDSIRISLLTCASG-GEIYSLFGHTAIRYENF-TRNIDAVFNYGM	75
Bfrag          ---------TESLSTADSIRISLLTCAPG-EEIYSLFGHTAIRYEEP-ARGIDRVYNYGL	71
Bfineg         ---------GPAADGHDSIRISLLTCAPG-EEIYSLFGHTAIRYENP-SQGIDVVFNYGL	64
Btheta         ---------QAIASGNDSIRLSLLTCAPG-EEIYSLFGHTAIRYEDP-ANGIDAVFNYGL	69
Bcaccae        ---------QTTANSNDSIRLSLLTCAPG-EVIYTLFGHTAIRYENP-SQGIDVVFNYGL	69
Bxylani        ---------QAPANSNDSIRLSLLTCAPG-EEIYSLFGHTAIRYENP-SQGIDAVFNYGL	69
Bovatus        ---------QASANNNDSIRLSLLTCAPG-EEIYSLFGHTAIRYENP-SQGIDIVFNYGL	69
Bnordii        ---------GTPDNNADSIRLSLLTCAPG-EEIYSYFGHTAIRYEEP-SKGIDWVFNYGI	70
Bsalyer        ---------DVFRNDNDSIRLSLLTCAPG-EEIYSYFGHTAIRYEDP-GKGIDVVFNYGL	69
Pplebeius      ----EISSDIKKVPQTDSIQFSLLTCAPG-SEIYALFGHTAIRYQNF-SKGVDLVFNYGM	79
Pmassi         -----ILASWSVRAQQDSVRVSLLTCAPG-TEIYELFGHTAIRYENP-AEGKDLVFNYGI	63
Pdorei         -----FSLSQSIRGQEDNIKVSLMTCAPG-TEIYALFGHTALRYEDT-ARGEDWVFNYGM	65
Bvulg          -----FFLGQSVRGQEDNIKVSLMTCAPG-TEIYALFGHTALRYEDK-ARGEDWVFNYGM	65
                                   ..::* .     ::  :**:::*  :      *   *:*

Osplan         FDFSTPHFYLKYARGLLPYQLTVQKFSHFIYNYQLEERTVYAQTIRLDSLEKQRIFDLLE	151
Barn_int       FNFDAPGFIWRFTKGETDYCLGINDFPDFLLNYQFRESKVDEQVLNLTPIQSRALFEALL	136
Pdista         FDFSKPNFIYRFAKGETDYKLGVADFQDYVIEYQMRGSDITEQVLNLTQEEKEHIWDALL	127
Pjohns         FDFSKPNFIYRFAKGETDYRLAAQYTRDFLIEYEMRGSEVTEQILDIDSAGKARIWEALM	128
Pmerdae        FDFSKPNFIYRFAKGETDYRLAAQYTRDFLIEYEMRGSEVTEQILDIDSAGKARIWEALM	128
Pgold          FDFSKPNFIYRFTKGETDYKLAAYNFSHYIIEYQMRGSEVTEQVLNLTPEETHKIWNALI	127
Pgord          FDFSKPNFIYRFTKGETDYKLAAYNFQHYIIEYQMRGSEVTEQVLNFTPDEINKIWNALY	127
Dgadei         FNMSKSNFIFLFMKGETDYMVAPIPYNTYYQEYKERGVGIIEQVFNLTQKEKQDIFDALL	132
Dmossii        FDFNSDNFIYRFVKGETYYMVGSIPFKYYIEEYGQRGVGVTEQVYNLTLKEKQEIFDALA	136
Pcopri         FSFNQSCFVLRFVFGLTDYQIGIYPMSDFIAEYAHEGRWVRQQRLRLSRTEKLGILRSID	153
Begger         FNFNAPNFILRFALGETDYQLGAGDYERFAAEYYYLERDVWQQELNLTPAEKKKLVTLLE	137
Bclarus        FNFNAPNFILRFALGETDYQLGVNSYERFAAEYHYLERDVWQQELNLTPQEKERLIALLE	135
Bsterc         FNFNAPNFILRFALGETDYQLGVTDYERFAAEYYYLERDVWQQELNLTVQEKEKLVMLLE	113
Bunif          FNFNAPNFIFRFALGETDYQLGVTDYEHFAAEYNYLGRDVWQQTLNLTEEEKERLITLLT	136
Bfluxus        FDFNTPNFTLRFALGDTDYQLGATNYRHFVSEYHSLGRDVWQQTLNLTQDEKERLINRLE	135
Boleici        FNFNAPNFIFRFALGETDYQLGVTSYERFAEEYDYLGRDVWQQTLNFTQDEKEHLVRLLQ	125
Bcell          FNFNAPNFIFRFALGETDYQLGVTNYEHFASEYNYLGRDVWQQTLNLTQAEKEHLFNLLQ	135
Bintesti       FNFNAPNFILRFALGETDYQLGATNYEHFVAEYNYLGRDVWQQTLNLTPDEKEHLFNLLQ	135
Bfrag          FSFNTPNFILRFALGKTDYQLGVEDYRRFAAEYEYFGRSVWQQTLNLTVEEQQQLITLLE	131
Bfineg         FSFNTPNFILRFSLGETDYQLGATDYAHFAAEYAFFGRSVWQQTLNLTEREKAELIRLLQ	124
Btheta         FSFNTPNFILRFSLGETDYQLGATDYARFAAEYAFDGRSVWQQTLNLSKEEKAELIRLLQ	129
Bcaccae        FSFDIPNFALRFSLGETDYRLGVIDYPHFAGEYAYFGRSVWQQTLNLNNEEKAELIRLLQ	129
Bxylani        FSFNTPNFIFRFSLGETDYQLGVTDYEHFAAEYAFYGRSVWQQTLNLTDEEKTKLIQLLQ	129
Bovatus        FSFNTPNFIFRFSLGETDYQLGATDYERFAAEYAFFGRSVWQQTLNLTDEEKTELIRLLQ	129
Bnordii        FNFGAPNFIFRFALGQTDYILGGMNYDRFAAEYIFDERSVWQQTLNLTPEEKQKLLALLI	130
Bsalyer        FNFGAPNFIFRFALGQTDYILGATPYNRFAAEYIFEERSVWQQTLNLTPDENRKLASLLI	129
Pplebeius      FSFDTPHFVYRFVKGETDYQLGITPYSYFESEYAFRGSSVYQQVLNLTYLEKISLLKLLQ	139
Pmassi         FSFNTPNFVLRFVKGETDYRLGVVPYSYFEGEYAVRGSSVYQQTLNLTDEEKQKIWDLLE	123
Pdorei         FSFNTPRFIYRFVKGETDYELGVTRYPYFEGSYAMRGSSVYQQTLNLTISEKQKLRRLLE	125
Bvulg          FSFNTPHFIYRFVKGETDYELGVTRYPYFEGSYAMRGSSVYQQTLNLTISEKQELRRLLE	125
               *.:.   *   :  *   * :       :  .*      :  *   :       :   :

Osplan         ENLLPQNRYYLYNFLFDNCTTRSRDILLKSLPDSV-------AWNMPDVN-KNFWNLLDE	203
Barn_int       VNAMPQNRVYRYNFLFDNCATRPRNMVEMVLDNKV-------RYKEPGESLPTFREEIDR	189
Pdista         INYRPENRVYRYNFFFDNCATRPAAILEKEINGSV-------DYQYPYQS-QTFRDLINY	179
Pjohns         INNRPENRVYRYNFFFDNCATRPAAIIENQTDGKI-------DYDAPFKQ-QTFRDLINY	180
Pmerdae        INNRPENRVYRYNFFFDNCATRPAAIIEKLAGGKI-------DYNVPFKQ-QTFRDLINY	180
Pgold          TNVQPENAVYRYNFFFDNCATRPVAIVEEQVDGKV-------QYNNPPEP-QTFRDLINY	179
Pgord          INVQPENCVYRYNFFFDNCATRPVAIVEEQVDGKI-------KYNDPPEP-QTFRDLINY	179
Dgadei         INCLPANREYRYNYFYDNCSTRPRDIFEKYINGKI-------EYTPTNKE-QTYRDLVIE	184
Dmossii        QNSNPENREYLYNFFYDNCATRPRDIVEKYVQGEI-------EYTPTDKQ-QTYRDLVWE	188
Pcopri         KNAQPENRVYRYNFFYDNCTTRAREMILSNLGNQS------TNFKDIPTA-STYREEIHK	206
Begger         ENYRPENRVYRYNFFYDNCATRPRDLIEKSIDGTL---QYADNMTDTNTG-TSFRDLLHK	193
Bclarus        ENYRPENRVYRYNFFYDNCATRPRDLIEKSIDGSL---QYADNMTDTNTG-TSFRDLLHK	191
Bsterc         ENYRPENRIYRYNFFYDNCATRPRDLIEKSINGTL---QYAGDMTDTDSG-ISFRDLLHK	169
Bunif          ENYRPENRVYRYNFFYDNCATRPRDQIERAINGTL---QYADNMTDNSTG-ISFRDLLHK	192
Bfluxus        ENYRPENRVYRYNFFYDNCATRPRDQIEKSIDGTL---QYADNMTDSNTG-VTYRDLLHK	191
Boleici        ENYRPENRVYRYNFFYDNCATRPRDQIESAIDGSL---QYTDNMTENNTG-VSFRDLLHK	181
Bcell          ENYRPENRIYRYNFFYDNCATRPRDQIEAAIDGTL---QYADNMTDTDTG-VTFRDLLHK	191
Bintesti       ENYRPENREYRYNFFYDNCATRPRDQIEKAIDGSL---QYADNMTDNNTG-VSFRDLLHK	191
Bfrag          ENYRPENRIYRYNFFYDNCATRPRDKVEESLQKSGSQLLFSNAHTENGET-KSYRDIVHQ	190
Bfineg         ENYRPENRVYRYNFFYDNCATRPRDKIEESIAGKV---IY--PAKPQDGS-RSFRDIVHQ	178
Btheta         ENYLPENRVYRYNFFYDNCATRPRDKIEESIDGKV---IY--PAEPQDGS-LSFRDIVHQ	183
Bcaccae        ENYRPENRVYRYNFFYDNCATRPRDKIEESIAGKV---IY--PVEPQDGS-RTFREIVHQ	183
Bxylani        ENYHPENRVYRYNFFYDNCATRPRDKIEESIAGKV---VY--PTEPQDGS-RTFRDIVHQ	183
Bovatus        ENYRPENRVYRYNFFYDNCATRPRDKIEESIAGKV---IY--PAEPQDGS-LTFRDIVHQ	183
Bnordii        ENSRPENRVYRYNFFYDNCATRPRDKIEESIQGKV---IY--NYPEKDGT-KSFRDIVHQ	184
Bsalyer        ENSKPENRTYRYNFFYDNCSTRPRDKIEECIEGKI---IY--DYPAKDGT-KSFREIVHQ	183
Pplebeius      DNYLPENRVYRYNYFYDNCTTRARDQIERSIQGKV-------VYPSVDWH-KTFRGIVHE	191
Pmassi         ENYRPGNRIYRYNYFYDNCTTRARDKIEDCIDGKV-------VYPQAEEG-VTFRDIVHR	175
Pdorei         ENYLPKNRVYRYNFFYDNCTTRARDIIEKCIEGKV-------VYSEGKER-LSFRDIVHQ	177
Bvulg          ENYLPENRVYRYNFFYDNCTTRARDVIERCIEGKV-------VYSEGKEG-LSFRDIVHQ	177
                *  * *  * **:::***:**    .                         .:   :

Osplan         YLQASPWVQWGIHTILGQRGNRTATTFQYMFLPDYLMYGLKNARYD-G----------QL	252
Barn_int       YAGICPWLIFGIDLALGSGLDRPMTYREQMFGPEILEKAFSEAVVQMS-------PDSAA	242
Pdista         CTRNHPWLTFGCDLALGSPTDREATQHEMLFLPPYLKEAFSKATIT-G-------PDGTI	231
Pjohns         CTRNRPWLTFGCDLALGSPTDRIATPHEMMFLPPYLKEAFGTATIT-G-------ADGSR	232
Pmerdae        CTRNKPWLTFGCDLALGSPTDRIATPHEMMFLPPYLKEAFGTATIT-G-------ADGSR	232
Pgold          CTRNHPWLTFGCDLALGSPTDRIATPHEMMFLPLYLKDEFEKATIV-N-------PDGSE	231
Pgord          CTRNNSWLTFGCDLALGSPTDRVATPHEMMFLPVYLKEEFDKATIV-N-------PDGTE	231
Dgadei         CTNSRQWFRFGINLVIGADADKVITDRQKDFLPRYLMNAYEGATIA-G-------D-SIP	235
Dmossii        CVGIQPWTKFGIDIIIGADADKVITDKQKDFLPAYLMKANEGARIK-N-------TDGTY	240
Pcopri         LNGQHRWARFGNDLLLGCQADRPITQREWEFLPDNLSKDFATEGRT-DFIDASQTSDGKT	265
Begger         YSKGHPWSRFGMDLCMGSQADKPISRRLMMFVPFYVQDYFNTARII-G-------SDKQV	245
Bclarus        YSEGHPWSRFGMDLCMGSQADEIINRRLMMFVPFYVQDYFNQAHIV-N-------KDGQA	243
Bsterc         YSKGHPWSRFGMDLCMGSQADKTINRRLMMFVPFYVQEYFNQARIV-N-------KEGET	221
Bunif          YSEGHLWSRFGMDLCMGSQADEPINRRLAMFVPFYMQEYFNKAQIV-D-------KEGQA	244
Bfluxus        YSEGHLWSRFGMDLCMGSKADKPISRREMMFIPFNVQEYFNTARIL-D-------KKGQA	243
Boleici        YSEGHPWSRFGMDMCMGSEADKPINRRLMMFIPFYVQEYFNTAQIV-N-------KEGQA	233
Bcell          YSEGHPWSRFGMDLCMGSKADQPINRRLMMFVPFYVQDFFNTARIV-D-------NEGQA	243
Bintesti       YSEGHPWSRFGMDMCMGSEADKPINRRLMMFVPFYVQEYFNTAQII-D-------KEGKA	243
Bfrag          YTKGHPWAQFGIDFCIGSQADHPINDRQMMFAPFYLMDAFAGARIA-N--------TSDN	241
Bfineg         YCKGHPWARFGIDLCIGSEADRPITQRQMMFAPFYLMDAFAGAQIE-N--------DSIQ	229
Btheta         YCKGHPWARFGIDLCIGSEADRPITQRQMMFAPFYLMDAFAGAQIV-H--------DSVQ	234
Bcaccae        YCKGHPWARFGIDLCIGGEADRPITQRQMMFAPFYLMDAFAGAQIT-G--------DSIQ	234
Bxylani        YCKGHPWARFGIDLCIGSEADQPITQRQMMFAPFYLMDAFDGAQIS-T--------DTYS	234
Bovatus        YCKGHPWARFGIDLCIGSEADQPITQRQMMFAPFYLMDAFDGAQIK-G--------DSIQ	234
Bnordii        YTNGHPWAQFGIDFCIGSEADRPITSRQMVFAPFYLKNALATAKIT-N--------NGNE	235
Bsalyer        YTQGHPWSQFGIDLCIGSEADRPITSRQMMFIPFYLEDAIASARIV-N--------NGNE	234
Pplebeius      FTKGSPWDELGIDLCLGAEADKPIDIRQQMFAPFYMRYFAQDAYIQ-T-------PDGVR	243
Pmassi         CTKGNEWDELGIDLCLGSEADVPIDGRKQMFAPFNMLEAARGAVIM-Q-------GDSV-	226
Pdorei         YTKGHEWDELGIDMCLGSEADNPIDTRKQMFAPFYMLEAAKKATIV-V-------GDSV-	228
Bvulg          YTKGHKWDELGIDMCLGSEADKPIDARKQMFAPFYLLEAAKKATIV-V-------GDSV-	228
                     *   * .  :*   :         * *  :

Osplan         -----------LAEPAEVLYQAPEMN----LSNSWYGTPLFVFALGCLLLILL--LQYVK	295
Barn_int       --------VPLVSRTEVLYDPEVPACPP---ETPFYLTPLFVAWLFFFFVAAVSVYDISR	291
Pdista         --------RPLVSETHVIGAGEADEPEK---DIWDLFTPLVCTWLLFGIVLGLTWIEWRK	280
Pjohns         --------KKLVSSTKTLVNGLADEER----PDTGFFTPLVSCWAFFLVVLAVTFIEWRR	280
Pmerdae        --------KELVSSTKTLINGLTDDVK----PDTGFFTPLVCCWAFFLVVLAVTFVEWRH	280
Pgold          --------RKLIKQTNRLAEQLTDDNN----GSKIWFTPMLCSMIFFLLIAVITFIEWKK	279
Pgord          --------RKLVKSTTRLAEELTDDDQ----GEKEWFTPMLCSMIIFLLVSLITYMEWKK	279
Dgadei         --------RNILLSTNTILEAKPFEKD-------FPVTPLYAGIILLIISILISYIVYKK	280
Dmossii        --------RNFVLKEQQLLMPQELVSE-------NHIQPLYVGCILLLITILLSFLVYKK	285
Pcopri         NATSASGYITLVDETSDIIPAQVQ------ITDDAPVTPQMIAIALAIIIIGTTVRECIK	319
Begger         --------RPLVLNEEKIITTGMEETGQ----PSEGFTPLQAALLLFILTAAATLYGIRR	293
Bclarus        --------RPLVLNEEKIIMTGSEEAEQ----PSDGFTPMQAALLLFILTAAATIYGIRR	291
Bsterc         --------RPLVLNEEKIIVTGNEEAEQ----PSDGFTPMQTALLLLILTAAATIYGIRR	269
Bunif          --------RPLVAKEEKIVVTGKTPADF----VSRGITPMQSASLLLILVAGISIYGIRR	292
Bfluxus        --------RPLVTSEEKVVITGKTAADF----QTGGISPMQSALLLFVIVTAATIYGIRK	291
Boleici        --------RPLVSAEEKIVVTGLTDADH----RSGGITPMQASLLLFIVVTAVTIYGIRH	281
Bcell          --------RPLVSSEEKIVVTGLTDADH----RSGGITPMQSALLLFVLVTAATIYGIRR	291
Bintesti       --------RPLISSEEKIVETGLTAADH----RSSGITPMQSALLLFILVTAATIYGIRR	291
Bfrag          --------KALVASTKKIIDCEPDVSDSAENDIWNMLPPIRLSLLVFIAIGMATVYGLRK	293
Bfineg         --------RPLAEAARLIVDATPGTDE-----SGWIFTPLQCALLLFILTVGATIYGIRR	276
Btheta         --------RPLVSGKELIVDALPEEEE-----GGWMPTPFQCSLLLFILTAAATIYSIRK	281
Bcaccae        --------RPLVTDSELIVDATPEEGE-----NFWIPTPLQSALLLFILTAAATIYGIRR	281
Bxylani        --------RPLVKTNELIIDVTPEPDE-----SGWMPTPLQCSLLLFILTAAATIYGIRR	281
Bovatus        --------RPLITANELVVDATPEPDE-----SDWMPTPLQCSLLLFILTAAATIYGIRR	281
Bnordii        --------RPLVADTELIINCEDENSSPAPISIADIFTPMRSALLLFIIVAGTSIYGIRK	287
Bsalyer        --------RPLVLESTSLISCEDDGSSPGTIGFWDLLTPIRAALLLLILTAAATIYGIRR	286
Pplebeius      --------RPLVLREEKIVDAELD-TE-----SSRKITPMIAAALFLLLNVIIGFFQWRK	289
Pmassi         --------RPLVLSESKVVDVEPEEAE-----PGFPLSPLSCVCILLVVTCIIVWLQLKW	273
Pdorei         --------RPLILHEKKVVDVEPEDVR-----EGFPLSPMVCVFILIGVTCFVGWLQFKI	275
Bvulg          --------RPLILHEKKVVDAEPENVG-----DGFPLSPLACVFILIGITCFVGWLQFKT	275
                                :                    *

Osplan         -SRSLLNVISLIFLLFTGIVGILIVFLGGFTAHPITAPNWNILWANPLNLFVIPFIFRKA	354
Barn_int       KRY--SRVFDTVLFSIYGLGGLVVFFLMFVSVHPATYPNYSAFWLHPFWLLMALFIWFKS	349
Pdista         KKY--FLWVDCVLFSVAGAGGVILFFLSFISVHPCTWPNWSLVWLQPLDLIAVILFCVKK	338
Pjohns         KSY--FRIVDCLLFLIAGIAGIVLFFLSFVSTHPCVCPNWNIIWLQPFDLAAVILFTVKK	338
Pmerdae        KSY--FRIVDCILFFTAGIAGIVLFFLSFVSTHPCVCPNWNIIWLQPFDLAAVILFTVKK	338
Pgold          KTY--FRMVDCILFFLAGLAGTVLFFLCFISTHPCIWPNWSIVWLQPFALIAVILFAVKK	337
Pgord          KTY--FRLVDCILFLLAGVAGTVLFFLCFISTHPCIWPNWSIVWLQPFDLVAVILFAVKK	337
Dgadei         KMVGLGKAFDTTLFLIAGIAGSIIFFLMFFSVHPCTNPNWNIIWLNPLQLIVVLLFYVKS	340
Dmossii        RWFTLGQIYDTLLFIVAGIGGCVIFFLMFFSIHPCVNPNWNIVWLNPLQLIVACLFFVKS	345
Pcopri         KKN--YWWFDAILLVLTGLPGLIL-FTMIFSQHPTVQINFQILILNPLNLIFAWKTVKRM	376
Begger         KKT--LWGIDLVLFFAAGMAGCILTFLVLFSQHPAVSPNYLLFVFHPLHLLCLPCMLNRV	351
Bclarus        KKT--LWGIDLALFFAAGTAGCILAFLALFSQHPAVSPNYLLFVFHPLHLFCLPCMLNRV	349
Bsterc         KKT--LWGIDFALFFAAGTAGCILAFLALFSQHPAVSPNYLLFVFHPLHLFCLPCMLNRV	327
Bunif          GKT--LWGMDLILFLVAGMAGCILAFLALFSQHPAVSPNYILFVFHPFHLFCLPFMLSRV	350
Bfluxus        GKT--LWGIDLILFFAAGVAGCILAFLALFSQHPAVSPNYLLFVFHPFHLLCLPCMLNRV	349
Boleici        QKT--LWGVDLLLFFCAGVAGCILAFLALFSQHPAVSPNYLLFVFHPLHLFCLPWMINKV	339
Bcell          GKT--LWGLDLILFFCAGIAGCILAFLALFSQHPAVSPNYLLFVFHPLHLFCLPWMINKV	349
Bintesti       RKT--LWGLDLILFFCAGIAGCILAFLALFSQHPAVSPNYLLFVFHPLHLFCLPWMINKV	349
Bfrag          KKS--LWGLDIAVFAAAGIAGCIIAFLALFSEHPTVGSNYLLFVFHPGHLLCLPFFINDE	351
Bfineg         RTG--LWGVDLFLFSAAGIVGCALAFLASFSEHPAVSSNFLLLVFHPGQLLFLPYIIYCD	334
Btheta         RTG--LWGVDLILFGAAGIVGCVLAFLALFSEHPAVSSNFLLLVFHPGQLLLLPYIIYCV	339
Bcaccae        RTG--LWGVDLILFGTAGIAGCILAFLALFSEHPAVSSNFLLFVFHPDQLLFLPYIIYCV	339
Bxylani        RTG--LWGIDLVLFGMAGIVGCVLAFLALFSQHPAVSSNFLLLVFHPGQLLFLPYIVYCV	339
Bovatus        RTG--LWGIDLFLFGIAGIVGCVLAFLALFSQHPAVSSNFLLLVFHPGQLLFLPYIIYCV	339
Bnordii        KRG--LWGIDLILFGIAGIAGCIIAFLALFSEHPAVSSNYLLFVFHPGELIFLPIIVNAA	345
Bsalyer        NKS--LWGIDIIWFGTAGIAGCILAFLALFSEHPTVSSNYLLFVFHPGQLVFLPIMINAA	344
Pplebeius      QMI--FWGWDILLYAVQGMAGCIIAFLFFISSHPTVGSNWLLLLFNPLPLFYLPFMIYRE	347
Pmassi         KKV--IWAWDLLLFGAQGLAGCIIAFLFFFSIHPTVGSNWLIVLFNPIPLIYLPVMIYKA	331
Pdorei         RKI--IWIWDLLLFGVQGLAGCVITFLVFFSTHPTVGSNWLILLLNPIPLIYLPVMVYRA	333
Bvulg          RKI--IWIWDLLLFGVQGLAGCVITFLVLFSTHPTVGSNWLILLLNPIPLLYLPVMVCRA	333
                        .       *  *  : *   .: **    *:  .  :*  *

Osplan         LPLFIRKYLKFY--LAILAIAIPVWAIAQPAVPMASISLIILM-IYLCFRLRNEK-----	406
Barn_int       LKSIVRYYHFAN--FAGLLLFVALWHWIPQQFNAAFFPLVLVLVIRSFTYLAVSV-----	402
Pdista         LKKAAYYYHFIN--FAALTLMLLGWHFIPQHLNSAFIPLVASIWVRSGWGVYRKI-----	391
Pjohns         LRKAAYYYHFIN--FAALTLMLAGWHFIPQHLNTAFIPLVMSIWLRSGYGVYRKI-----	391
Pmerdae        LRKAAYYYHFIN--FAALTLMLAGWHFIPQHLNTAFIPLVMSLWLRSGYGVYRKI-----	391
Pgold          FGKAAYYYHFIN--FAALTLLLAGWYFIPQHLNIAFIPLVATLWLRSGYGIYRGN-----	390
Pgord          YGKAAYYYHFIN--FAALTLLLAGWYFIPQHLNIAFIPLVATLWLRSGFGVYRGR-----	390
Dgadei         LSKCIYYYHFIN--FVALLAFLLAWNLIPQQLEVTFIPFILSIGLRSFMNILQQK-----	393
Dmossii        LSKFISCYHFIN--FVALLAFLLAWCLIPQQLEIAFIPFILSLCIRSGMNILQLK-----	398
Pcopri         KAGRQYWYFELLGWLLLIALFMQIW----QNYAEGMSILALTLLARYCVKSTMMDLSPNN	432
Begger         RKRRRSRYMLAN--FLVLTLFILLWLIIPQRFPSAVLPLALCLLIRSASNL---------	400
Bclarus        RKRKRSRYMLAN--FIVLTLFILLWLVIPQRFPLAVLPLALCLLIRSASNL---------	398
Bsterc         RKRKRSRYMLAN--FIVLTLFILLWLVIPQRFPLAVLPLALCLLIRSASNL---------	376
Bunif          RKREISRYMLVN--FTVLTLFIGLWAIIPQRIPPAILPLALCLLIRSASNL---------	399
Bfluxus        RKKKTSRYMIAD--LVVLTLFICFWAIIPQRIDLAVLPLALCLLIRSASNL---------	398
Boleici        RKRQKSRYMVAN--FIVLTLFILLWAIIPQRFDLAVLPLALCLLVRSASNL---------	388
Bcell          RKRQKSWYMRTN--CAILTLFILLWAIIPQRIDLAVLPLALCLLVRSASNL---------	398
Bintesti       RKRQKSRYMVLN--FIVLTLFILLWAIIPQRFDLAVLPLALCLLVRSASNL---------	398
Bfrag          RKRRKSRYHLLN--YTVLTLFIVLFPVIPQNFDLAVLPLALCLLIRSASNL---------	400
Bfineg         RKGKKCWYLTLN--LIVLTLFIVLFPLIPQRFDLAVVPLALSLLVRSVSNL---------	383
Btheta         RKGKKCWYLTLN--LVVLTLFMVLFPLIPQRFDLAVVPLALCLLIRSASNL---------	388
Bcaccae        RKGKKCWYLTLN--LIILTLFIVLFPVIPQRFDFAVVPLALCLLIRSVSNL---------	388
Bxylani        RKGKKCWYLTLN--LVVLTLFIVLFPVIPQRFDFAVVPLALVLLIRSASNL---------	388
Bovatus        RKGKKCWYLTLN--LAVLTLFIVLFPVIPQRFDFAVVPLALVLLIRSASNL---------	388
Bnordii        RKGRKCWYHMLN--CIVLTLFILLFPVIPQRIDLAVVPLALSLLIRSASNL---------	394
Bsalyer        RKGRKCWYHVLN--FTVLTLFILLFPLIPQRIDLAVVPLALSLLIRSASNL---------	393
Pplebeius      LKHKKDLYHTCN--LVYLTLFMVLFPFLPQDFNLTVLPLALGLLVNAASHV---------	396
Pmassi         IKGRKDLYHLIN--MAYLTFFIMIMPFIQQEFNVTVLPLALCLLVCSTGHV---------	380
Pdorei         IKGKKDYYHTIN--IVCLTSFMMIMPFIQQKFNVTVLPLALCLLICSANHV---------	382
Bvulg          IKGKKDLYHTIN--VAYLTLFIMIMPFVQQKFNVTVLPLALCLLICSANHV---------	382
                      *         :   :                :   :

Osplan         -----------------R--	407
Barn_int       ------RLKKTEKNKDEK--	414
Pdista         ------WRIG----------	395
Pjohns         ------WNIGYGKY------	399
Pmerdae        ------WNIGYEKY------	399
Pgold          ------MAIKGK--------	396
Pgord          ------MHI-----------	393
Dgadei         ------KFKKKADYSLPRAK	407
Dmossii        ------KLKKRADYSLPRRK	412
Pcopri         YGLQKHIVKKFRKNK-----	447
Begger         -------ILTYDKK------	407
Bclarus        -------ILTYDKK------	405
Bsterc         -------ILTYDKK------	383
Bunif          -------ILTYKRK------	406
Bfluxus        -------ILTYKRPS-----	406
Boleici        -------VLTSKKREKR---	398
Bcell          -------ILTLKKR------	405
Bintesti       -------ILTSKKR------	405
Bfrag          -------ILTYKKAK-----	408
Bfineg         -------IVTSKKK------	390
Btheta         -------ILTSKKK------	395
Bcaccae        -------IVTSKKK------	395
Bxylani        -------IVTSKKK------	395
Bovatus        -------IVTSKKK------	395
Bnordii        -------ILTYKKTK-----	402
Bsalyer        -------ILTYKKKK-----	401
Pplebeius      -------LVLNKK-------	402
Pmassi         -------FLYYRQNSK----	389
Pdorei         -------LLYYRQNNK----	391
Bvulg          -------LLYYRQNNK----	391
```

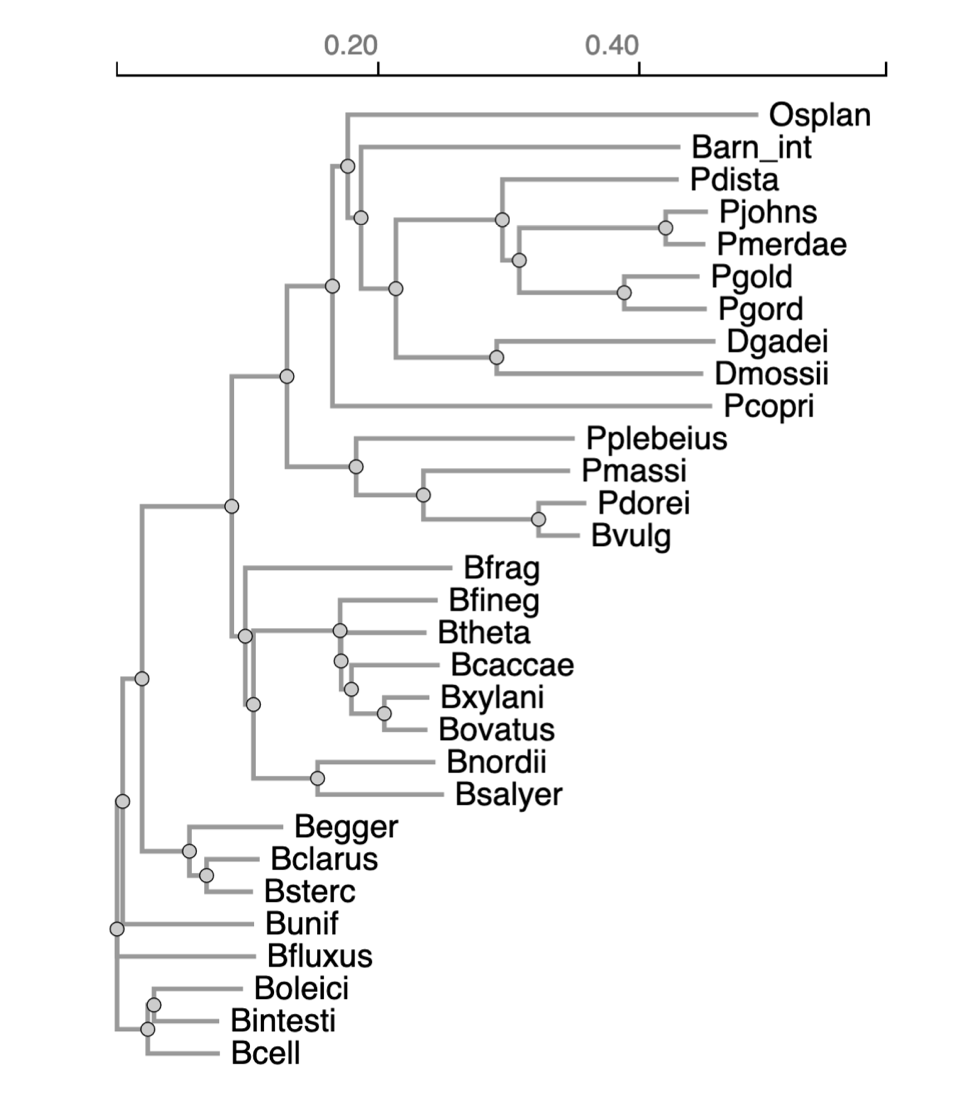
B.

C.

| **Species** | **Accession Number** | **Abbreviation (panels A&B)** |
| --- | --- | --- |
| *Bacteroides thetaiotaomicron* VPI-5482 | WP_008764530.1 | Btheta |
| *Bacteroides nordii* DSM 18764 | WP_025866461.1 | Bnordii |
| *Bacteroides fragilis* NCTC 9343 | WP_008768021.1 | Bfrag |
| *Bacteroides finegoldii* DSM 17565 | WP_032839999.1 | Bfineg |
| *Bacteroides xylanisolvens* DSM 18836 | WP_211421276.1 | Bxylani |
| *Bacteroides salyersiae* DSM 18765 | WP_055295141.1 | Bsalyer |
| *Bacteroides caccae* ATCC 43185 | WP_005680205.1 | Bcaccae |
| *Bacteroides oleiciplenus* DSM 22535 | WP_009128909.1 | Boleici |
| *Bacteroides cellulosilyticus* DSM 14838 | WP_195507924.1 | Bcell |
| *Bacteroides fluxus* DSM 22534 | WP_039968753.1 | Bfluxus |
| *Bacteroides intestinalis* DSM 17393 | WP_007666217.1 | Bintesti |
| *Bacteroides clarus* DSM 22519 | WP_195501148.1 | Bclarus |
| *Bacteroides stercoris* ATCC 43183 | WP_005657803.1 | Bsterc |
| *Parabacteroides goldsteinii* DSM 19448 | WP_224265698.1 | Pgold |
| *Phocaeicola plebeius* DSM 17135 | WP_007561568.1 | Pplebeius |
| *Parabacteroides johnsonii* DSM 18315 | WP_087375515.1 | Pjohns |
| *Phocaeicola dorei* DSM 17855 | WP_038607879.1 | Pdorei |
| *Phocaeicola massiliensis* DSM 17679 | WP_005936771.1 | Pmassi |
| *Bacteroides ovatus* ATCC 8483 | WP_195358720.1 | Bovatus |
| *Bacteroides uniformis* ATCC 8492 | WP_151875347.1 | Bunif |
| *Bacteroides eggerthii* DSM 20697 | WP_270569872.1 | Begger |
| *Dysgonomonas gadei* ATCC BAA-286 | 651385561 | Dgadei |
| *Bacteroides vulgatus* ATCC 8482 | 640762680 | Bvulg |
| *Dysgonomonas mossii* DSM 22836 | 651391713 | Dmossi |
| *Odoribacter splanicnicus* DSM 20712 | 649960002 | Osplan |
| *Barnesiella intestinihominis* DSM 21032 | 2530576248 | Barn_int |
| *Parabacteroides distasonis* ATCC 8503 | 2870967840 | Pdista |
| *Parabacteroides merdae* ATCC 43184 | 641042905 | Pmerdae |
| *Parabacteroides johnsonii* DSM 18315 | 2524924991 | Pjohns |
| *Prevotella copri* DSM 18205 | 2563257723 | Pcopri |

Fig S2. Lnb is conserved among gut Bacteroidota. (A) Multi-sequence alignment of Lnb homologs across 30 common type strains within the Bacteroidota, as used by Pudlo et. al. (1). The residues that comprise the putative catalytic triad Asn66-His49-Cys172 and Cys198 bolded in the *B. thetaiotaomicron* VPI-5482 sequence. Alignment created with CLUSTAL OMEGA tool (2). (B) Simple Phylogram from alignment within CLUSTAL OMEGA. (C) Table of sequences used for (A) and (B).

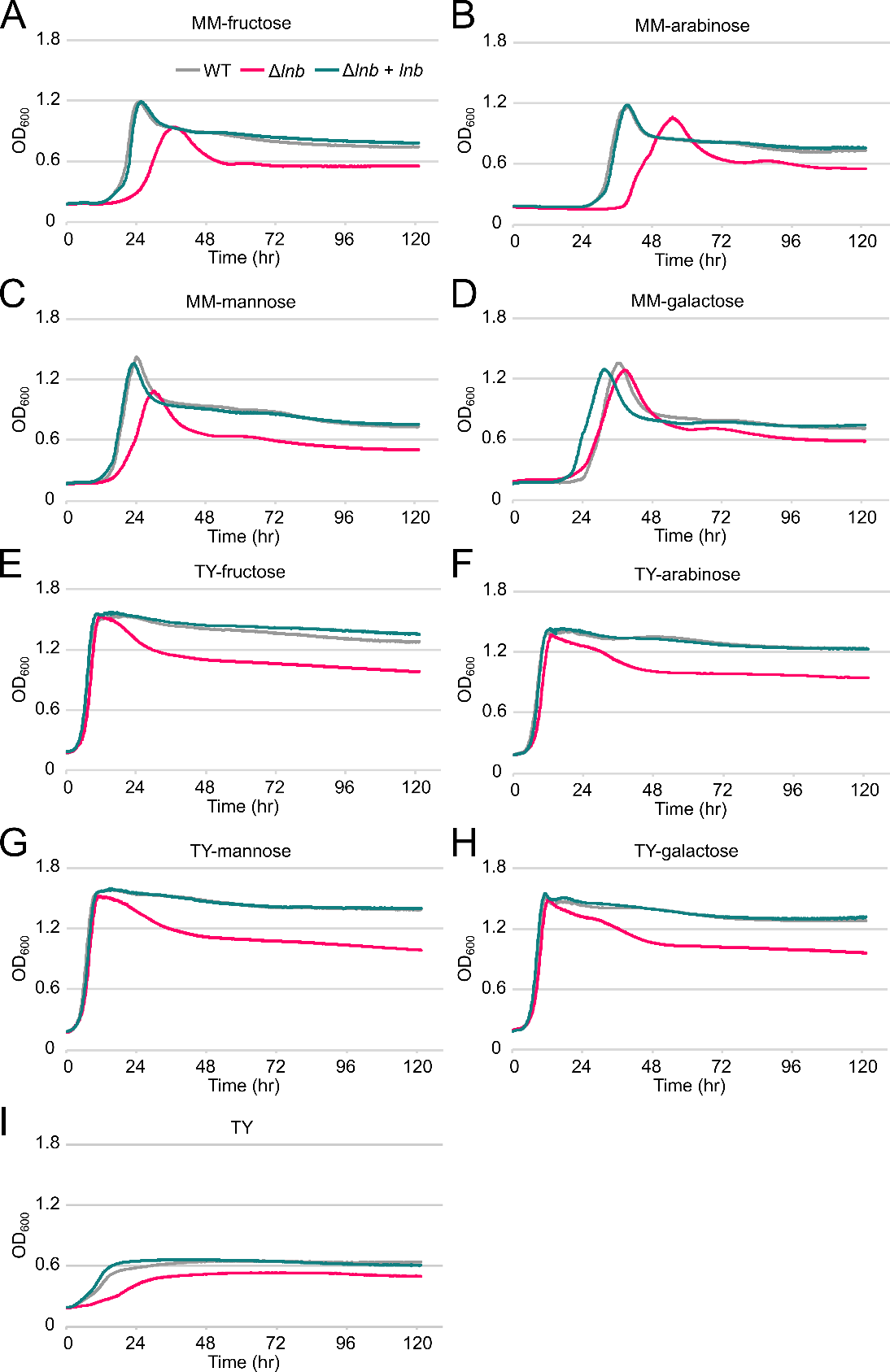

Fig S3. The optical density at 600 nm (OD_600_) of *B. theta* WT, Δ*lnb*, and complemented strains on MM-fructose (A), MM-arabinose (B), MM-mannose (C), and MM-galactose (D), TY-fructose (E), TY-arabinose (F), TY-mannose (G), TY-galactose (H), and TY with no additional carbon source (I) over time. Curves are the average of three technical replicates. The data shown is representative of three biological replicates.

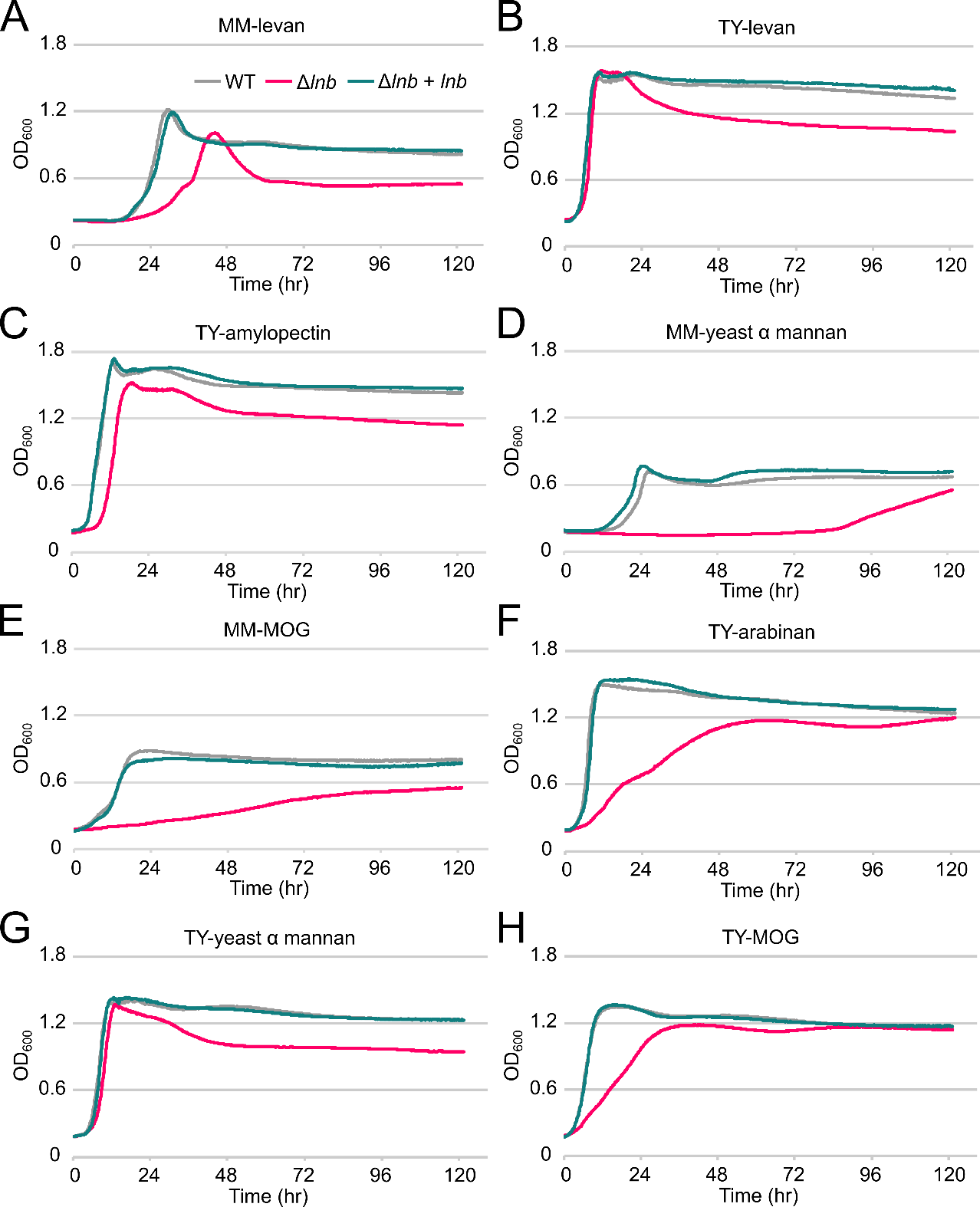

Fig S4. The optical density at 600 nm (OD_600_) of *B. theta* WT, Δ*lnb*, and complemented strains on MM-levan (A), TY-levan (B), TY-amylopectin (C), and MM-yeast α mannan (D), MM-MOG (E), TY-arabinan (F), TY-yeast α mannan (G), and TY-MOG (H) over time. Curves are the average of three technical replicates. The data shown is representative of three biological replicates.

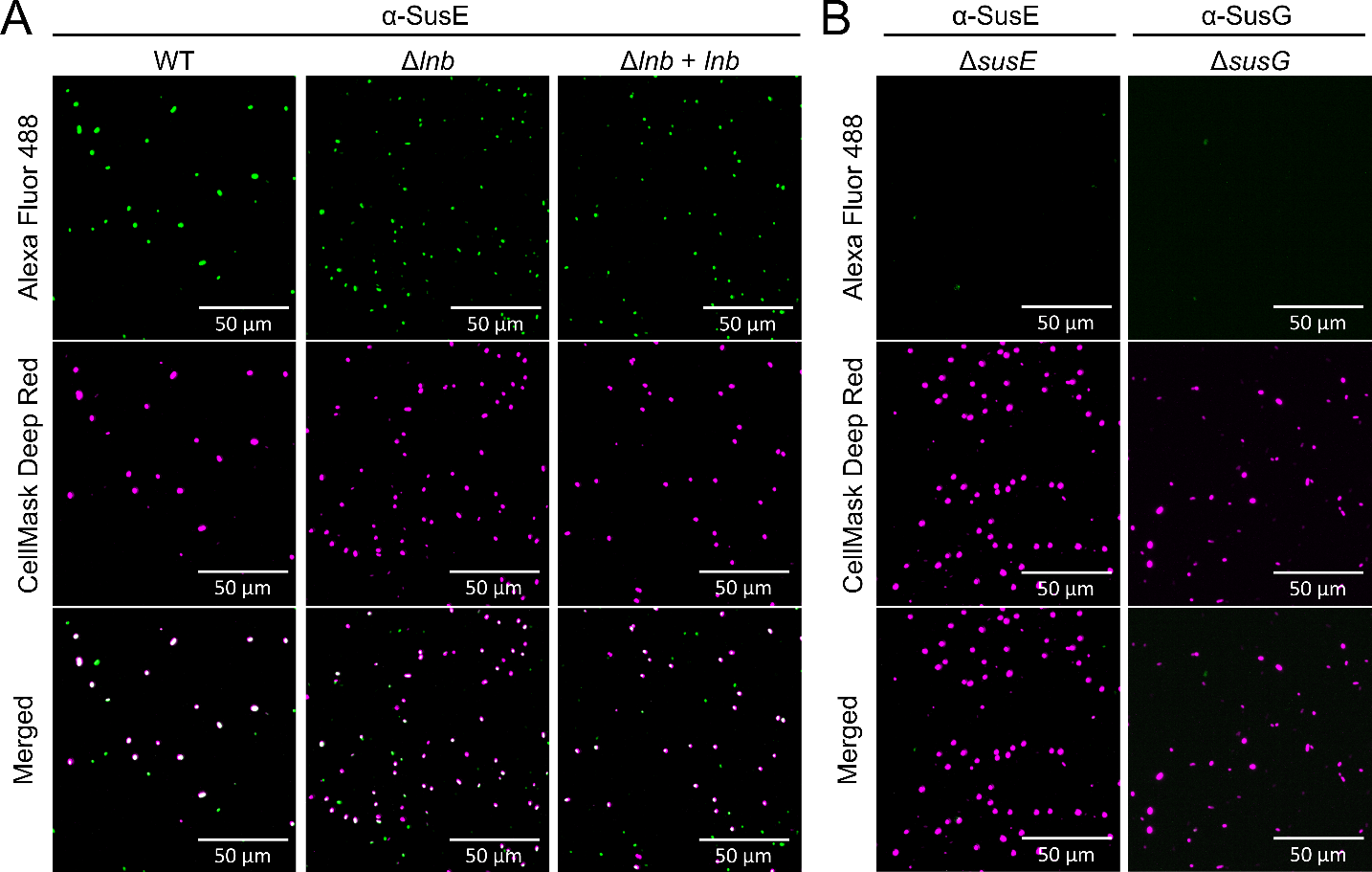

Fig S5. (A) Representative microscopy images of *B. theta* WT, Δ*lnb*, and complemented cells when immunostained for SusE. (B) Representative microscopy images of Δ*susE* cells immunostained for SusE and Δ*susG* cells immunostained for SusG. Alexa Fluor 488 and CellMask Deep Red images were taken in the green and red channels, respectively.

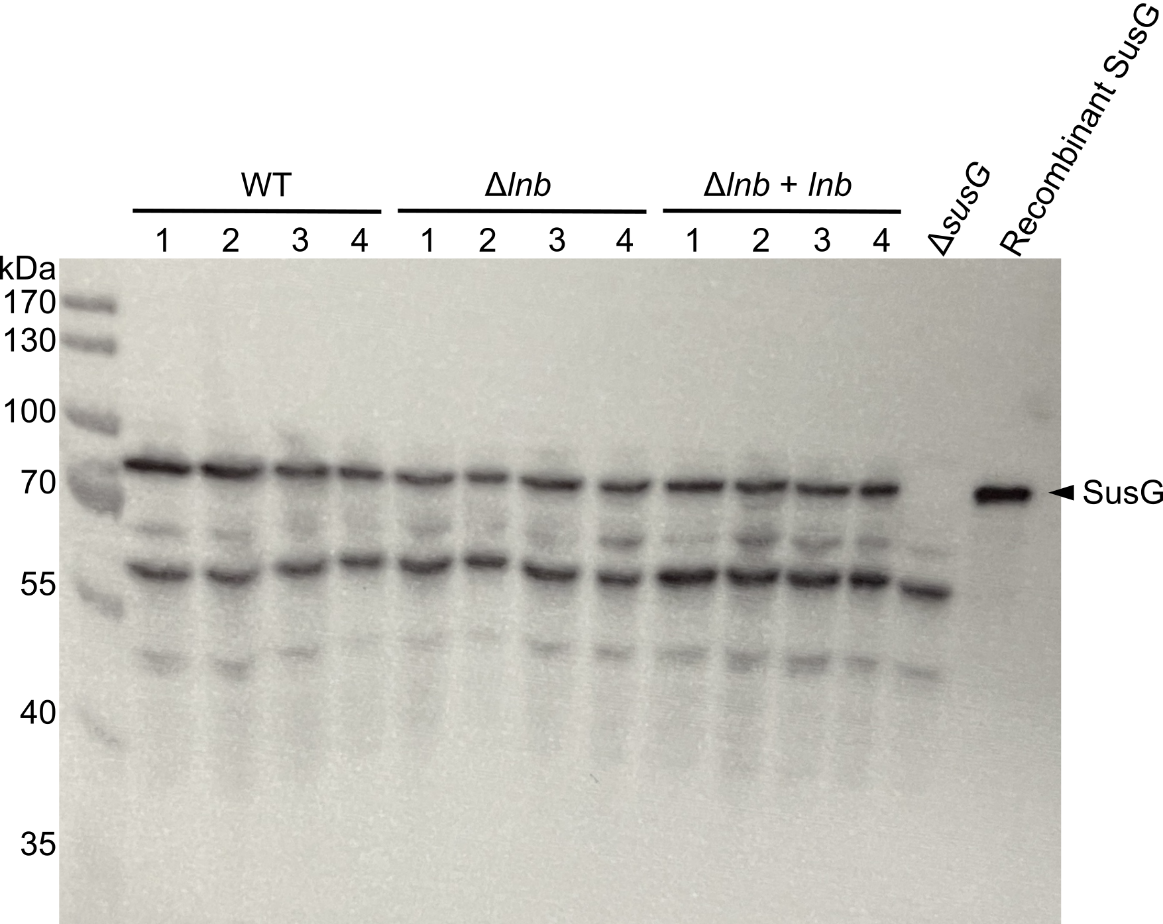

Fig S6. An immunoblot against SusG using whole-cell lysates of *B. theta* WT, Δ*lnb*, and complemented strains. Lysate of Δ*susG* cells and purified recombinant SusG protein are also included. The band for SusG is indicated by a black arrow. Cells were normalized by OD_600_.

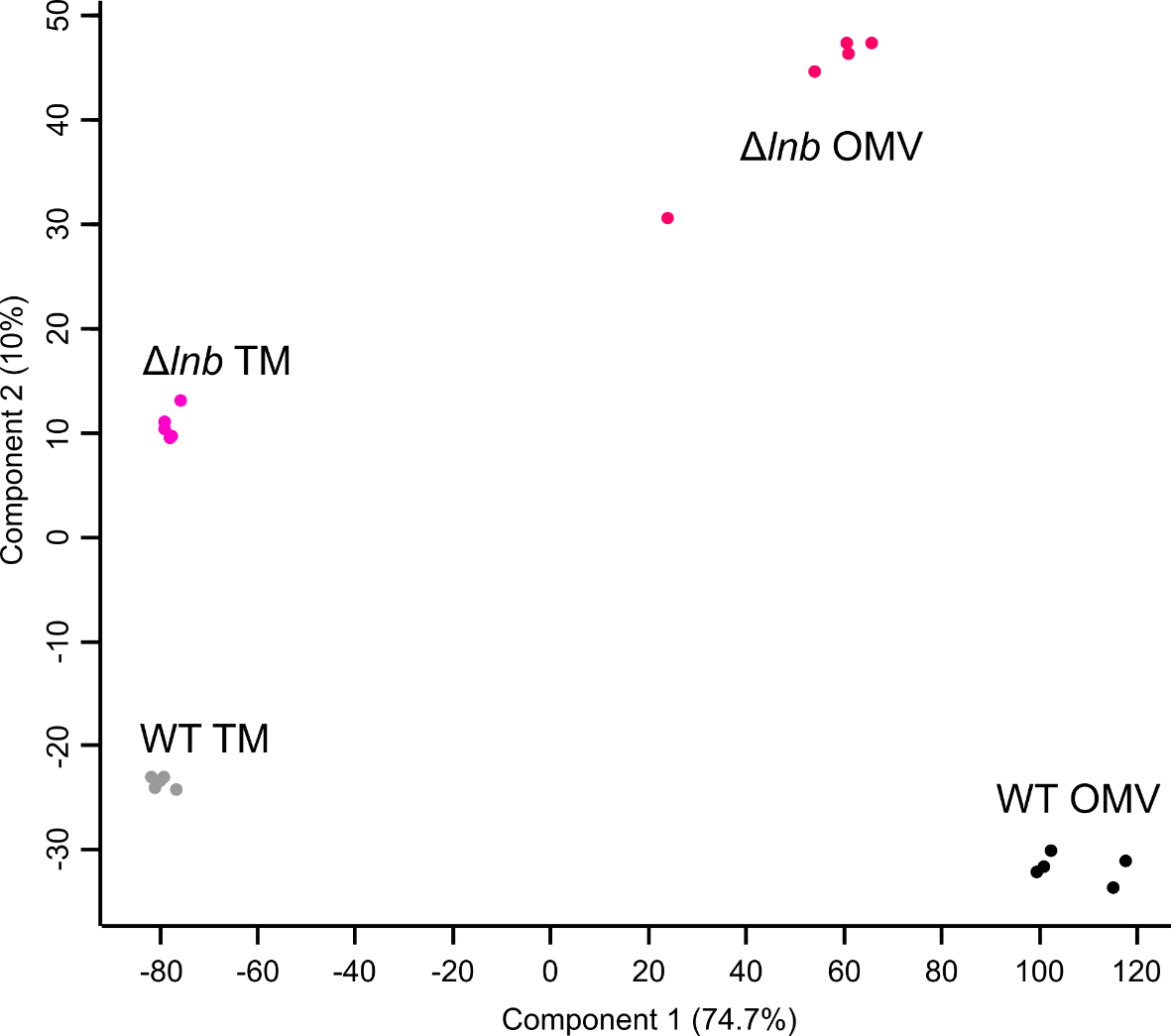

Fig S7. Principal component analysis (PCA) of total membranes and vesicles proteomes from *B. theta* WT (gray and black, respectively) and Δ*lnb* (magenta and red, respectively) grown in minimal media supplemented with starch. Five biological replicates were performed for each condition.

| **WT**  **(Total = 311)** | **Δ*lnb***  **(Total = 277)** | **Functional Prediction** |
| --- | --- | --- |
| 71 | 89 | Function unknown |
| 48 | 8 | Translation, ribosomal structure and biogenesis |
| 38 | 16 | Inorganic ion transport and metabolism |
| 23 | 14 | Signal transduction mechanisms |
| 21 | 10 | Energy production and conversion |
| 19 | 29 | Cell wall/membrane/envelope biogenesis |
| 19 | 21 | Carbohydrate transport and metabolism |
| 17 | 9 | Coenzyme transport and metabolism |
| 13 | 8 | Amino acid transport and metabolism |
| 8 | 21 | Transcription |
| 7 | 5 | Nucleotide transport and metabolism |
| 6 | 8 | Intracellular trafficking, secretion and vesicular transport |
| 5 | 11 | Defense mechanisms |
| 5 | 5 | Posttranslational modification, protein turnover, chaperones |
| 4 | 6 | Lipid transport and metabolism |
| 3 | 7 | Replication, recombination and repair |
| 3 | 3 | Cell motility |
| 1 | 3 | Cell cycle control, cell division, chromosome partitioning |
| 0 | 4 | Secondary metabolites biosynthesis, transport and catabolism |

Table S2. Functional analysis of proteins upregulated in total membranes from *B. theta* WT and Δ*lnb*. Proteins significantly enriched in the total membranes of each strain were analyzed using eggNOG-mapper.

Table S3. Strains and plasmids used in this study.

| **Strain or Plasmid** | **Relevant Genotype/Phenotype^a^** | **Reference** |
| --- | --- | --- |
| *Bacteroides fragilis* NCTC 9343 |  |  |
| *Bacteroides thetaiotaomicron* VPI-5482^b^ | Δ*tdk* |  |
| KA126 | pWW1376-*E. coli Lpp(K58A)-Strep* | This study |
| KA198 | Δ*BT_4364* | This study |
| KA201 | Δ*BT_4364* + pWW1376-*us1311-BT_4364* | This study |
| KA204 | Δ*BT_4364* + pWW1376-*E. coli Lpp(K58A)-Strep* | This study |
| KA200 | Δ*BT_4364* + pWW1376-*E. coli Lpp(K58A)-Strep-us1311-BT_4364* | This study |
| KA243 | Δ*BT_4364* + pWW1376-*E. coli Lpp(K58A)-Strep-us1311-BT_4364(C148S)* | This study |
| KA244 | Δ*BT_4364* + pWW1376-*E. coli Lpp(K58A)-Strep-us1311-BT_4364(C198S)* | This study |
| KA259 | pWW1376-*BT_3736-Strep* | This study |
| KA260 | Δ*BT_4364* + pWW1376-*BT_3736-Strep* | This study |
| KA263 | Δ*BT_4364* + pWW1376-*BT_3736-Strep-us1311-BT_4364* | This study |
| WT-tag | pNBU2-barcode 1 | This study |
| dLnb-tag | Δ*BT_4364* + pNBU2-barcode 14 | This study |
|  | Δ*susG* | (3) |
|  | Δ*susE* | (4) |
| Stellar *E. coli* | General cloning strain (Takara Bio) |  |
| *E. coli* S17-1 | *pir^+^* |  |
| *E. coli* BW25113^b^ | *E. coli* K-12 wild type; Δ(*araD araB*)*567* Δ*lacZ4787*(::*rrnB-3*) λ^-^ *rph-1*  Δ(*rhaD-rhaB*)*568 hsdR514* | CGSC7636^c^ |
| TXM327 | *lpp::*Chl^r^ | (5) |
| KA349 | TXM327 *ybeX-(*Kan^r^*-rrnB* TT*-araC-P_BAD_)-lnt* | (5) |
| TXM1036 | *fadR*::Tmp^r^ *lpp*::FRT *fadE*::Tet^r^ plus pTXM1026 [pBBR1(ori) *P_Kan_-lolCDE-PA3286* Kan^r^] *lnt*::Spt^r^ | (6) |
| KA98 | *lpp::*Chl^r^, *ybeX-(*Kan^r^*-rrnB* TT*-araC-P_BAD_)-lnt* + pKA27 + pUC19 | This study |
| KA118 | *lpp::*Chl^r^ *lnt*::Spt^r^ + rescuing plasmid p1 + pKA117 | This study |
| KA119 | *lpp::*Chl^r^ *lnt*::Spt^r^ + pUC19-*BF9343_0945* + pKA117 | This study |
| KA120 | *lpp::*Chl^r^ *lnt*::Spt^r^ + rescuing plasmid p31 + pKA117 | This study |
| Plasmids |  |  |
| pUC19 | General cloning vector; Car^r^ |  |
| pKA524 | pCL25-*E. coli Lpp(K58A)-Strep*, Spt^r^ | (5) |
| pKA117 | pCL25-*E. coli Lpp(K58A)-Strep*, Kan^r^ | This study |
| pExchange-*tdk* | Gene deletion in *Bacteroides*; Car^r^ in *E. coli*, Ery^r^ in *Bacteroides* |  |
| pWW1376 | Gene overexpression in *Bacteroides*; Kan^r^ in *E. coli*, Ery^r^ in *Bacteroides* | (7) |
|  | pNBU2-barcode 1; Car^r^ in *E. coli*, Tet^r^ in *Bacteroides* | (8) |
|  | pNBU2-barcode 14; Car^r^ in *E. coli*, Tet^r^ in *Bacteroides* | (8) |

^a^Resistance phenotypes: Chl^r^, chloramphenicol; Kan^r^, kanamycin; Spt^r^, spectinomycin; Car^r^, carbenicillin; Ery^r^, erythromycin; Tet^r^, tetracycline.
^b^Strains in gray boxes are derivatives of *Bacteroides thetaiotaomicron* VPI-5482 Δ*tdk* or *E. coli* BW25113.
^c^Strain CGSC7636 at the Coli Genetic Stock Center (CGSC).

Table S4. Primers used in this study.

| **Primer Name** | **Description** | **Primer Sequence** |
| --- | --- | --- |
| KA151 | 5'- pExchange *BT_4364* upstream | GAAGATAACATTCGAGCAGGATATTTATGCTTCATAGC |
| KA152 | pExchange *BT_4364* upstream -3' | GATAAAGAGTCAGTAATGG |
| KA156 | 5'- pExchange *BT_4364* downstream | CCATTACTGACTCTTTATCGCTCCTATTACCGATCTTCG |
| KA159 | pExchange *BT_4364* downstream -3' | GGTGGCGGCCGCTCTAGGTCAACCTCATCGACGATG |
| KA116 | 5'- pUC19 *BF9343_0945* | CGGTACCCGGGGATCCGCCACATGAACTCAAAGTTG |
| KA117 | pUC19 *BF9343_0945* -3' | CCATGATTACGCCAAGCTTGTACAGTCAGTATGGAGG |
| KA118 | 5'- pUC19 *BF9343_0946* | CGGTACCCGGGGATCCCAGAGAGAAAGTGAAC |
| KA119 | pUC19 *BF9343_0946* -3' | CCATGATTACGCCAAGCTTGAAGCGCACAAAGTGTC |
| KA120 | 5'- pUC19 *BF9343_0947* | CGGTACCCGGGGATCCCTATAATAATGGTAGG |
| KA121 | pUC19 *BF9343_0947* -3' | CCATGATTACGCCAAGCTTATACCGTCTATTCTCTC |
| KA168 | 5'- pUC19 *BF9343_0944* | CGGTACCCGGGGATCCGCAATCTTATTCTGACATAC |
| KA169 | pUC19 *BF9343_0944* -3' | CCATGATTACGCCAAGCTTTGCGCTTTGCACCTTG |
| KA177 | 5'- *BT_4364* upcheck | CTAATCAGAGGCTCTGTTTGGG |
| KA178 | *BT_4364* downcheck -3' | GATTGTCTGCACGGTGC |
| KA219 | 5'- pWW1376 inverse | GCTATTACGAGCGCTTAAACGG |
| KA220 | pWW1376 inverse -3' | CGGTATCAAGTCATAGCAGTC |
| KA221 | 5'- pWW1376 *us1311* | GACTGCTATGACTTGATACCGTGATCTGGAAGAAGC |
| KA222 | pWW1376 *us1311 BT_4364* -3' | CCGTTTAAGCGCTCGTAATAGCATTATTTCTTTTTAGAGGTC |
| KA191 | 5'- pWW1376 *E. coli Lpp* | CAATAATTTATTTTCAATGAAAGCTACTAAACTGG |
| KA192 | 5'- *us1311* NotI | CCCTCCACCGCGGTGGCGGCCGCTGATCTGGAAGAAGC |
| KA193 | *us1311* NdeI -3' | GTTCTAGATAGTGCCATATGTTAAAAACAGATTTGG |
| KA179 | 5'- pWW1376 Strep inverse | GGATCCGGTTCAGCGTGGAGTCATCCTCAATTTGAAAAATAAGGTTCCTAGCTGATTAG |
| KA85 | EcLpp Strep-3' | CAAATTGAGGATGACTCCACGCTGAACCGGATCCCG |
| KA194 | 5'- *us1311 BT_4364* | CAAATCTGTTTTTAACATATGAAACGGACAATTC |
| KA282 | 5'- *BT_4364* C198S | CATCGACCTCAGCATCGGTAG |
| KA283 | *BT_4364* C198S -3' | CTACCGATGCTGAGGTCGATG |
| KA284 | 5'- *BT_4364* C148S | CTTTTATGACAACAGCGCTACACGCC |
| KA285 | *BT_4364* C148S -3' | GGCGTGTAGCGCTGTTGTCATAAAAG |
| KA136 | 5'- pKA524 *kanR* | GTACCTGTGAAGTGAAGCTTCAAATATGTATCCGCTCATG |
| KA137 | pKA524 *kanR* -3' | CGCAGGGGATCAAGATCTCTGTCTGCTTACATAAACAG |
|  | 5'- barcode 1 | ATGTCGCCAATTGTCACTTTCTCA |
|  | 5'- barcode 14 | GGCACGCCATTCTTCATCTAACTG |
|  | universal barcode -3' | CACAATATGAGCAACAAGGAATCC |
| KA372 | 5'- pWW1376 *BT_3736* | CAATAATTTATTTTCAATGAAAAAATTATTATTTGCG |
| KA373 | pWW1376 *BT_3736* Strep -3' | CCACGCTGAACCGGATCCGTCAAGTTTTTTACTTAC |
| KA378 | pWW1376 Strep -3' | CAGCTAGGAACCTTATTTTTCAAATTGAGGATGACTCCACGCTGAACCGGATCC |
